# The FANCM-RMI1/2 complex promotes genomic instability and PARP inhibitor sensitivity in BRCA2-deficicient cells

**DOI:** 10.64898/2026.01.15.699681

**Authors:** François Dossin, Mikael Attia, Fabricio Montero Zúñiga, Daniel González Acosta, Rachel Brough, Fei Fei Song, Aurélien Boré, Raphaël Margueron, Massimo Lopes, Christopher J. Lord, Ludovic Deriano

## Abstract

Cancer cells lacking BRCA1 or BRCA2 must adapt to survive and proliferate despite defective DNA repair and high genomic instability. Restoration of homologous recombination (HR) through loss of DNA end protection rescues growth defects and promotes PARP inhibitor (PARPi) resistance in BRCA1-deficient cells; however, the genetic basis of adaptation to BRCA2 loss remains largely unexplored. To delineate BRCA1- and BRCA2-specific adaptation trajectories, we established a fully isogenic screening platform in mouse embryonic stem cells engineered for acute depletion of either protein. This approach uncovered hundreds of genes with shared, distinct, or even opposing effects between the two BRCA-deficiency states. We identified FANCM and its interacting partners RMI1/2, CENPS/MHF1, CENPX/MHF2, and FAAP24 as essential in BRCA1-deficient cells but toxic in BRCA2-deficient contexts. Loss of FANCM in BRCA2-deficient cells alleviated genomic instability and proliferation defects by preventing RMI1/2-dependent replication fork degradation, without rescuing HR. Moreover, we show that the FANCM–RMI1/2 complex drives PARPi sensitivity in both mouse and human BRCA2-deficient cells, in contrast to the 53BP1–SHLD–CST axis in BRCA1-deficient cells. These findings reveal distinct adaptation routes to BRCA1 and BRCA2 loss and establish FANCM as a determinant of BRCA2-specific vulnerability and therapeutic response, with direct implications for tailoring prevention and therapy according to BRCA mutation status.

## INTRODUCTION

Loss-of-function mutations in the tumor suppressor genes *BRCA1* and *BRCA2* are associated with breast, ovarian, prostate and pancreatic cancers. However, unlike in cancer cells, where it provides proliferative advantage, loss of *BRCA1*/*2* in healthy cells results in significant fitness costs *in vitro* and early embryonic lethality in mice ^1–3^. This is likely due to the vital role BRCA1/2 play in maintaining genome integrity and the deleterious impact of aberrantly resolved DNA lesions that occurs upon their loss. How cancer cells adapt to survive and proliferate despite the loss of BRCA1/2 and increased genome instability is not fully understood.

BRCA1 and BRCA2 are essential for repairing DNA double strand breaks (DSB) through homologous recombination (HR) and for protecting stalled replication forks against nascent strand degradation ^4,5^. However, despite being involved in similar processes, BRCA1 and BRCA2 are mechanistically distinct proteins. For instance, BRCA1, but not BRCA2, promotes DSB end resection during the initial stages of HR ^4,5^ and also controls DSB repair by single strand annealing (SSA), a BRCA2-independent process ^6^. Furthermore, BRCA2 (and its partner PALB2) is directly involved in loading the RAD51-recombinase onto single-stranded DNA, while BRCA1 only plays an indirect role, and suboptimal RAD51 loading can occur in the absence of BRCA1 ^7^. Consistent with these mechanistic differences, BRCA1- and BRCA2-mutated tumors differ in their histopathology, genomic signature and mechanism of resistance to therapy ^8–11^. These observations suggest that adaptation trajectories could potentially differ between *BRCA1*- and *BRCA2*-mutated cancers. Unravelling the biological basis of this adaptation is essential for anticipating the emergence of *BRCA1*- and *BRCA2*-mutated cancer cells in patients, developing tailored treatments, and predicting how tumors may resist therapy. In this study, we aimed to systematically identify the genetic routes through which healthy noncancerous cells adapt to the loss of BRCA1/2, and in turn determine whether these routes differ in BRCA1- as opposed to BRCA2-deficient contexts. To understand this, we generated a purely isogenic BRCA-proficient (non-adapted) cellular system in which either BRCA1 or BRCA2 can be conditionally inactivated. To this end, we used wild-type mouse embryonic stem cells (mESCs) in which we conducted parallel genome-wide CRISPR-screens before inducing acute degradation of BRCA1 or BRCA2 using the auxin-inducible degron system (AID2) ^12^.

## RESULTS

We first engineered a parental mES cell line amenable to both CRISPR-screening and auxin-inducible degron (AID2) by stably integrating CAS9 and OsTIR1^F74G^ constitutive expression cassettes at the safe-harbor *Tigre* locus (**Figure S1A**). We then derived two isogenic cell lines, hereafter referred to as BRCA1-degron (BRCA1^AID^) and BRCA2-degron (BRCA2^AID^), in which an AID-GFP cassette was homozygously knocked-in and translationally fused to the N-terminal coding sequences of endogenous BRCA1 or BRCA2, respectively (**Figure 1A and Figure S1B**).

**Figure 1:**
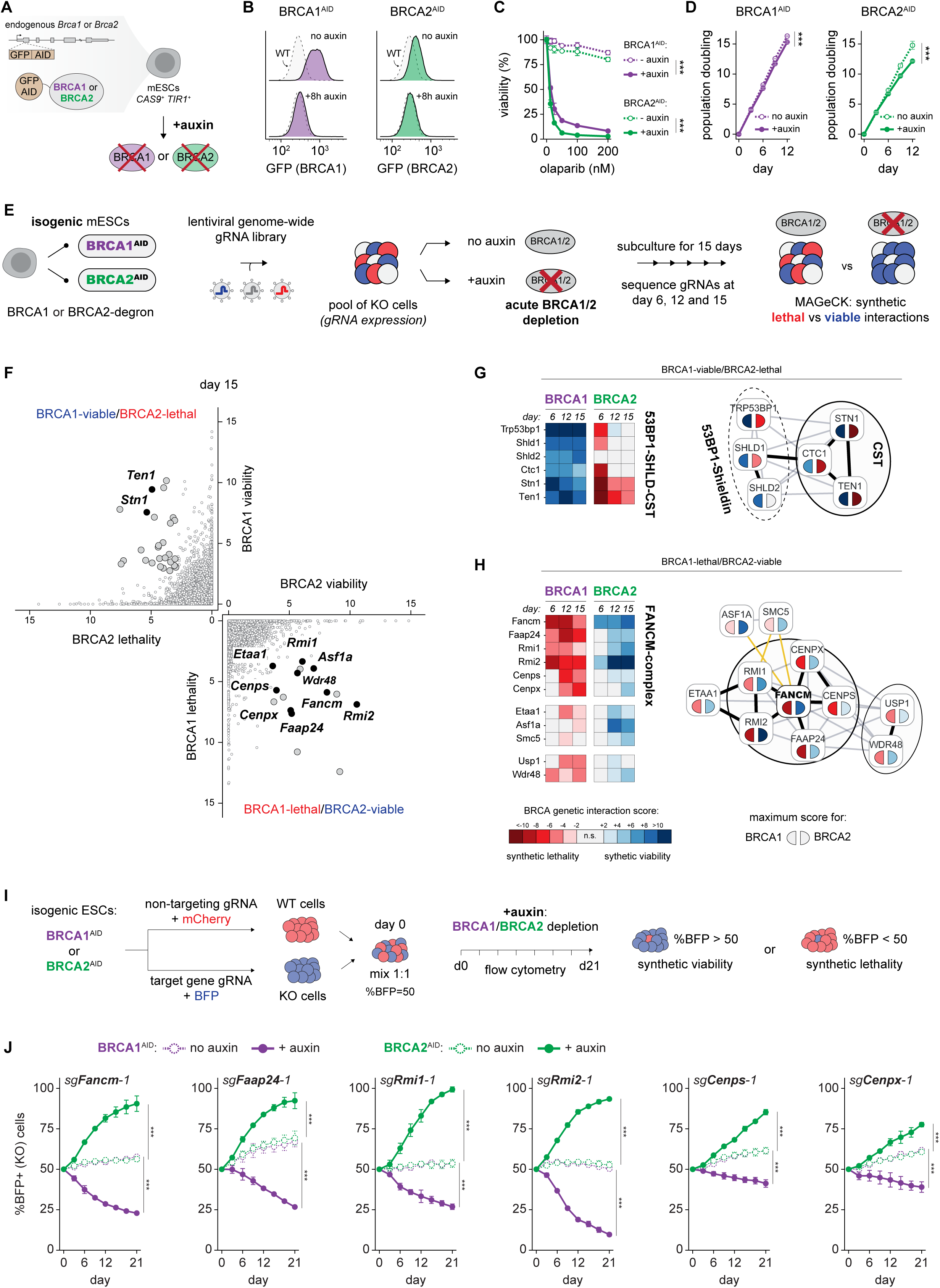
Isogenic BRCA degron screening identifies opposite genetic interactions between BRCA1- and BRCA2-deficient cells. **A**, Schematic of isogenic BRCA1 and BRCA2-degron mESCs. **B**, Flow cytometry analysis of auxin-induced degradation of endogenous GFP-tagged BRCA1 and BRCA2. **C**, Cell survival after 5 days of BRCA1 or BRCA2 degradation (+auxin) and exposure to increasing concentrations of olaparib. **D**, Cumulative population doubling over a 12-day long time course of BRCA1 or BRCA2 depletion (+auxin). **C** and **D**, represent the mean ±SD of 2 biological replicates. Statistical analysis: two-way ANOVA; ***P<0.001. **E**, Schematic of the CRISPR-screen in isogenic healthy BRCA-degron mESCs to identify synthetically lethal and viable interactions with acute BRCA1 or BRCA2 loss. **F**, Scatterplot showing BRCA1-SV versus BRCA2-SL scores (top-left quadrant) and BRCA1-SL versus BRCA2-SV scores (bottom-right quadrant) at day 15. Genes meeting significance threshold (FDR<0.05 and RRA<0.05) are represented as larger dots. **G** and **H**, Heatmaps showing synthetic lethal (red shades) and viable (blue shades) interaction scores (MAGeCK RRA) for different genes at the three timepoints of the BRCA1- (left heatmap) and BRCA2- (right heatmap) screens. STRING interaction networks for each gene groups are shown on the right, with genes colored according to their maximal BRCA-interaction score in each screen. **I**, Schematic of the two-color competitive fitness assays to validate screen hits. **J**, Competitive fitness assays during a 21-day time course of BRCA1- (purple) or BRCA2- (green) depletion, monitoring BFP-positive cells expressing gRNA-pairs targeting FANCM and its physical interactors. The mean ±SD of 2 independent experiments is shown. Statistical analysis: two-way ANOVA; ***P<0.001.

Auxin treatment resulted in rapid and efficient depletion of BRCA1 and BRCA2 protein (**Figure 1B**) and expectedly rendered cells exquisitely sensitive to the PARP inhibitor olaparib (**Figure 1C**). In the absence of exogenous genotoxic stress, the acute loss of BRCA1 or BRCA2 over a 12-day long period reduced cell proliferation, particularly in BRCA2-depleted cells, but did not result in complete lethality (**Figure 1D**). Importantly, this sensitized context enables the simultaneous identification of positive (hereafter termed synthetic viable or SV) and negative (hereafter termed synthetic lethal or SL) genetic interactions ^13^, involving genes whose loss would improve or worsen the fitness of BRCA-depleted cells, respectively.

To systematically identify such interactions, we performed parallel genome-wide CRISPR screens upon auxin-mediated BRCA1 or BRCA2-depletion in the isogenic BRCA-degron cells (**Figure 1E**). Cells were collected at different timepoints of auxin-exposure (**Figure 1E**) to capture a wide dynamic range of BRCA1/2-related genetic interactions, anticipating that certain sensitive interactions might be specific to either short or long periods of BRCA-depletion. MAGeCK analysis yielded a cumulative total of ∼950 SV and ∼660 SL interactions across the BRCA1 and BRCA2 screens (**Figure S1C** and **Table S1**), with marked differences observed between early and late depletion timepoints (**Figure S1E, F** and **Table S1**). Reactome analysis of these significant genes revealed a strong enrichment for DNA repair pathway terms, particularly DNA repair, DSB repair and HR-related ones (**Figure S1D** and **Table S2**), demonstrating the sensitivity and specificity of our screens in uncovering biologically meaningful genetic interactions.

The intrinsic isogenicity of the BRCA1/2-degron cells (**Figure 1A, E**) allows for direct comparison of the BRCA1- and BRCA2-centered genetic interaction networks. Interestingly, only a small fraction of synthetic lethal and synthetic viable genes are shared by both networks (**Fig S1C**). Shared SL interactions include DNA repair pathways such as base excision repair (BER) and non-homologous end-joining (NHEJ), as well as several unexpected chromatin regulatory complexes (see **Supplementary text** and **Fig S2** for further information). Shared SV interactions, involving genes whose loss improves the fitness of both BRCA1- and BRCA2-deficient cells, include P53 as well as the anti-resection factors DYNLL1 and HELB (see **Supplementary text** and **Fig S2)**. Conversely, our approach identified several genetic interactions which are specific to either the BRCA1- or the BRCA2-deficient context. Notably, loss of RAD51-paralogs, the BRCA1-A/C complexes and EXO1 were synthetic lethal with BRCA1-loss but not BRCA2-loss, while loss of POLQ was, conversely, synthetic lethal with BRCA2-loss but not BRCA1-loss (see **Supplementary text** and **Fig S2)**.

We next interrogated our dataset to determine whether a given gene could simultaneously be involved in opposite genetic interaction with BRCA1 and BRCA2. In other words, the loss of such a gene should reduce the fitness of BRCA1-deficient cells while increasing that of BRCA2-deficient cells, or vice versa. The existence of such gene would be of significant clinical importance because its perturbation would have antagonistic outcomes (*i.e.* cell fitness cost versus benefit) depending on BRCA1- versus BRCA2-mutation status. We therefore filtered for genes that simultaneously met significance thresholds for synthetic viability with BRCA1 and synthetic lethality with BRCA2 (or vice versa) at any of the three screen time points. Remarkably, forty-two BRCA1-viable/BRCA2-lethal and twenty BRCA1-lethal/BRCA2-viable genes meeting those criteria were identified (**Figure 1F and Figure S3A, B, Table S3**).

Among the BRCA1-viable/BRCA2-lethal genes, components of the CST complex (*Ctc1*, *Stn1* and *Ten1*) showed robust antagonistic genetic interactions with BRCA1 (synthetic viability) and BRCA2 (synthetic lethality), at all time points (**Figure 1F, G and Figure S3A, B**). Regarding BRCA1, 53BP1 (*Trp53bp1*) and subunits of the Shieldin complex (*Shld1* and *Shld2*) behaved similarly to CST, being synthetic viable throughout the screen (**Figure 1G**). However, with respect to BRCA2, 53BP1-Shieldin showed either mild synthetic lethality, viability, or no significant genetic interaction depending on the timepoint (**Figure 1G** and **Figure S3A**). We set out to clarify these findings using an orthogonal, candidate-focused approach. To this end, we used a two-color fitness assay in which the relative growth of candidate-KO cells (BFP-positive) in competition against control cells (mCherry-positive) is monitored during a 21-day time course of BRCA1 or BRCA2 depletion in the isogenic BRCA-degron lines (**Figure 1I**). In this assay, gRNAs targeting 53BP1 (*Trp53bp1*), Shieldin (*Shld1*, *Shld2*) and CST (*Ctc1*, *Stn1*, *Ten1*) all conferred a proliferative advantage to BRCA1-depleted cells (**Figure S3C**). 53BP1-Shieldin removal also improved the growth of BRCA2-depleted cells, albeit to a lesser extent than in BRCA1-depleted cells (**Figure S3C**, see discussion). Conversely, loss of all CST subunits resulted in strong synergistic growth defects with BRCA2 depletion (**Figure S3C**). Taken together, our results demonstrate that CST is essential for the survival of BRCA2-deficient cells specifically, through a function that is independent of 53BP1-Shieldin. One such function could be CST’s role in protecting stalled replication forks from nucleolytic degradation in BRCA2-deficient cells ^14^. Thus, while loss of CST benefits the growth of BRCA1-deficient cells and promotes PARPi resistance in BRCA1-mutated cancer ^15,16^, we argue that inhibiting CST function would in fact be detrimental to the survival of BRCA2-mutated cancers and might represent a promising therapeutic avenue.

Next, we focused on antagonistic genes whose loss is synthetically lethal with BRCA1-loss but viable with BRCA2-loss. This analysis identified the USP1-WDR48 complex (**Figure 1F, H and Figure S3A, B**), which deubiquitinates PCNA and FANCD2 (see discussion). Most strikingly, the BRCA1-lethal/BRCA2-viable analysis revealed the DNA translocase FANCM at all time points, as well as its known direct physical interactors: RMI1, RMI2, FAAP24, CENPS (MHF1) and CENPX (MHF2) (**Figure 1F, H and Figure S3A, B**). Other antagonistic genes included ASF1A and SMC5, which interact physically with the yeast ortholog of FANCM (Mph1 ^17–19);^ and ETAA1 which interacts physically with RMI1/2 ^20^ (**Figure 1F, H and Figure S3A, B**). In validation assays, we confirmed that sgRNAs targeting *Fancm* and its direct partners *Rmi1*, *Rmi2*, *Faap24*, *Cenps* and *Cenpx* all result in synergic growth defects upon depletion of BRCA1 but conversely provide proliferative advantage to BRCA2-depleted cells (**Figure 1J** and **Figure S3D**). Therefore, FANCM appears to be the central hub of a physical network whose loss yields opposite fitness outcomes in BRCA1- versus BRCA2-compromised cells, which we decided to investigate in more detail.

We first focused on characterizing the synthetic viability between *Fancm* and *Brca2* by generating independent *Fancm*-KO clones in the BRCA2-degron background using CRISPR-CAS9 gene targeting (**Figure S4A, B**). Consistent with a direct role of FANCM in the repair of interstrand crosslinks (ICLs) by the Fanconi anemia (FA) pathway ^21^, *Fancm*-KO clones were hypersensitive to the ICL-inducing agent mitomycin C (MMC, **Figure S4C**). We first assessed the durability of the *Fancm*/*Brca2* synthetic viability by monitoring population doubling times of *Fancm*-WT and *Fancm*-KO cells during four weeks of BRCA2-depletion. In *Fancm*-WT cells, BRCA2-removal strongly reduced proliferation and increased doubling time by 5-hours (**Figure 2A, B and Figure S4D**). BRCA2-depleted *Fancm*-WT cells were enriched in G_1_- and depleted in S-phases of the cell cycle (**Figure S4E**), consistent with previous reports ^22,23^. Remarkably, loss of FANCM restored growth throughout the 4-week period of BRCA2-depletion, to a level similar to that of BRCA2-proficient cells (**Figure 2A, B and Figure S4D**). Cell cycle alterations associated with BRCA2-deficiency were also rescued in *Fancm*-KO cells (**Figure S4E**). These results are consistent with the idea that healthy cells can durably adapt to the loss of BRCA2 by inactivating *Fancm*.

**Figure 2:**
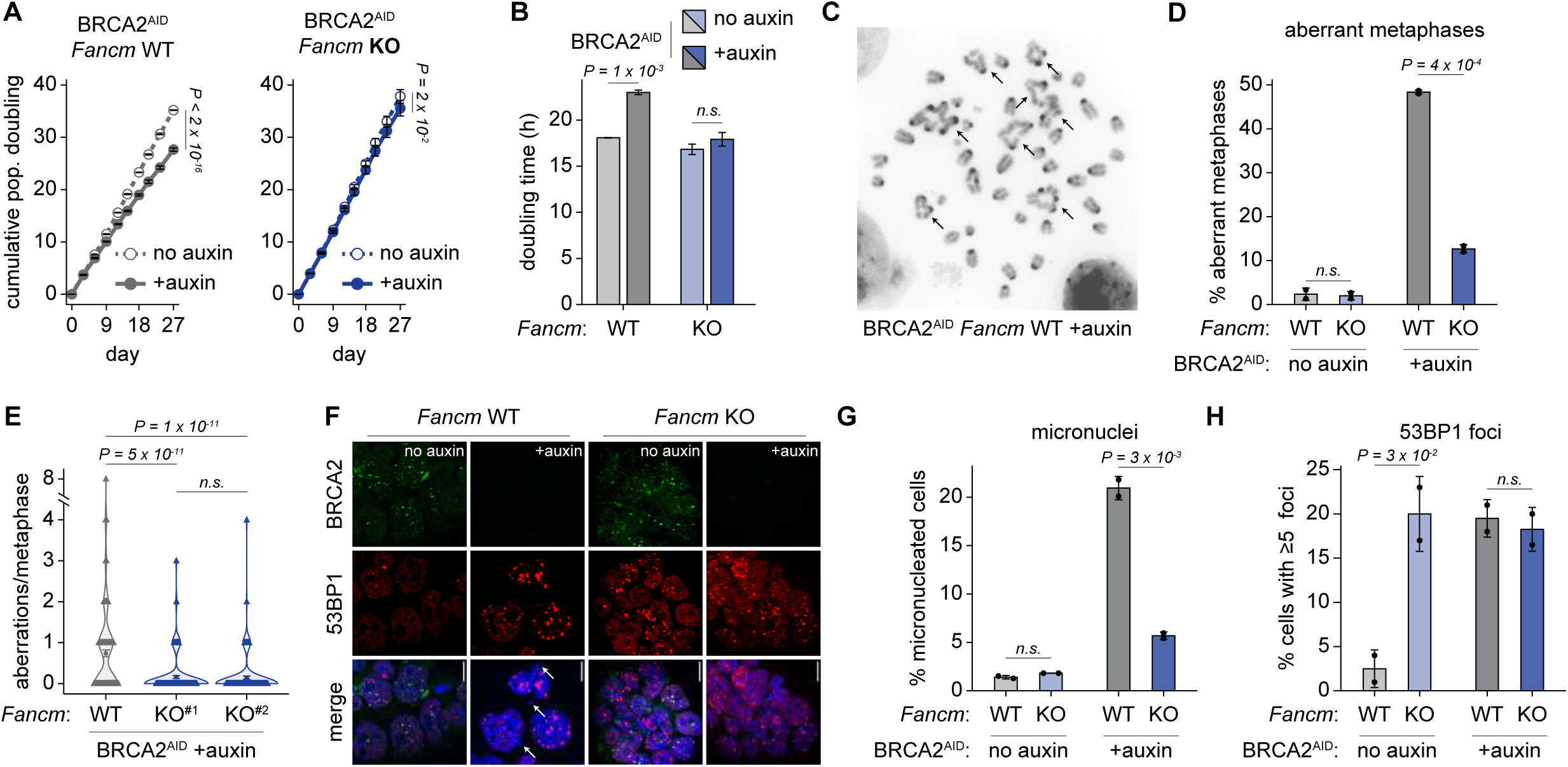
Loss of FANCM restores proliferation and genome stability in BRCA2-deficient cells. **A**, Cumulative population doubling and time of *Fancm*-WT and *Fancm*-KO cells during 27 days of BRCA2-depletion (+auxin). The mean ±SD of 2 independent clones is shown. Statistical analysis: two-way ANOVA. **B**, Population doubling times computed from Fig. 2a. Statistical analysis: two-sided Student’s t-test. **C**, Metaphase spread following 5 days of BRCA2-depletion in *Fancm*-WT cells. Arrows indicate chromosome aberrations. **D**, Barplots representing the fraction of cells with at least one aberrant metaphase following 5 days of BRCA2-depletion in *Fancm*-WT or *Fancm*-KO cells. The mean ±SD of 2 independent clones is shown. n=150 metaphases were scored. Statistical analysis: two-sided Student’s t-test. **E**, Violin plots representing the number of chromosomal aberrations per cell among metaphases scored in Fig. 2d. The mean ±SEM of 150 metaphases is shown as a circle. Statistical analysis: two-sided Wilcoxon rank-sum test. **F**, Representative images of micronuclei (white arrows) and 53BP1 foci, **G**, barplots representing the fraction of cells with micronuclei or **H**, more than five 53BP1 foci in *Fancm*-WT and *Fancm*-KO cells upon BRCA2-depletion for 5 days (+auxin). Data in **G** and **H**, represent the mean ±SD of 2 independent clones, with n≥100 cells scored for micronuclei/53BP1 foci. Statistical analysis: two-sided Student’s or Welch’s t-test.

Next, we investigated whether *Fancm*-removal rescues the genomic instability associated with BRCA2-deficiency by monitoring chromosomal abnormalities, spontaneous micronuclei formation, and 53BP1 foci (a hallmark of DNA break-sites) during 5 days of auxin-mediated BRCA2-depletion. In *Fancm*-WT cells, BRCA2-depletion resulted in a large increase in aberrant metaphases (close to 50% of cells carried at least one chromosome abnormality), micronuclei and 53BP1 foci formation (**Figure 2C-H**). However, these defects were largely rescued in *Fancm*-KO cells, with only 10% of cells harboring chromosome abnormalities (**Figure 2D-H**). Importantly, BRCA2 nuclear signal (including BRCA2-foci) was undetectable following auxin treatment in both *Fancm*-WT and KO cells (**Figure 2F**), ruling out the possibility that loss of *Fancm* would artifactually promote viability through abrogation of AID-dependent BRCA2-degradation. It is worth noting that while loss of FANCM had no effect on the chromosomal integrity of BRCA2-proficient cells (**Figure 2D, G and Figure S4F**), 53BP1 foci were nevertheless higher in *Fancm*-KO than in *Fancm*-WT cells (**Figure 2F, H**), consistent with FANCM’s role in limiting 53BP1-foci formation ^24^. However, in Fancm-KO cells, 53BP1 foci levels remained unchanged upon BRCA2-depletion, in contrast to *Fancm*-WT cells (**Figure 2H**).

Collectively, our data show that FANCM promotes genomic instability in BRCA2-deficient cells, and that its loss protects BRCA2-deficient cells from such instability and consequently increases their proliferative capacity.

To gain insight into how FANCM exerts its toxicity in BRCA2-deficient cells, we first conducted parallel genome-wide CRISPR screens in *Fancm*-WT and *Fancm*-KO cells, in both BRCA2-proficient (no auxin) and BRCA2-deficient (+auxin) contexts (**Figure 3A**). In BRCA2-proficient conditions, *Smarcal1* was the most significantly depleted gene in *Fancm*-KO cells compared to *Fancm*-WT cells (**Figure 3B**), consistent with recent reports of a very strong synthetic lethal interaction between *FANCM* and *SMARCAL1* ^25,26^. Furthermore, *Brca1* and its partner *Bard1* were also identified as synthetic lethal hits in the *Fancm*-KO screen (**Figure 3B**), in line with the *Fancm/Brca1* genetic interaction uncovered reciprocally in the BRCA1-degron screen (see **Figure 1**).

**Figure 3:**
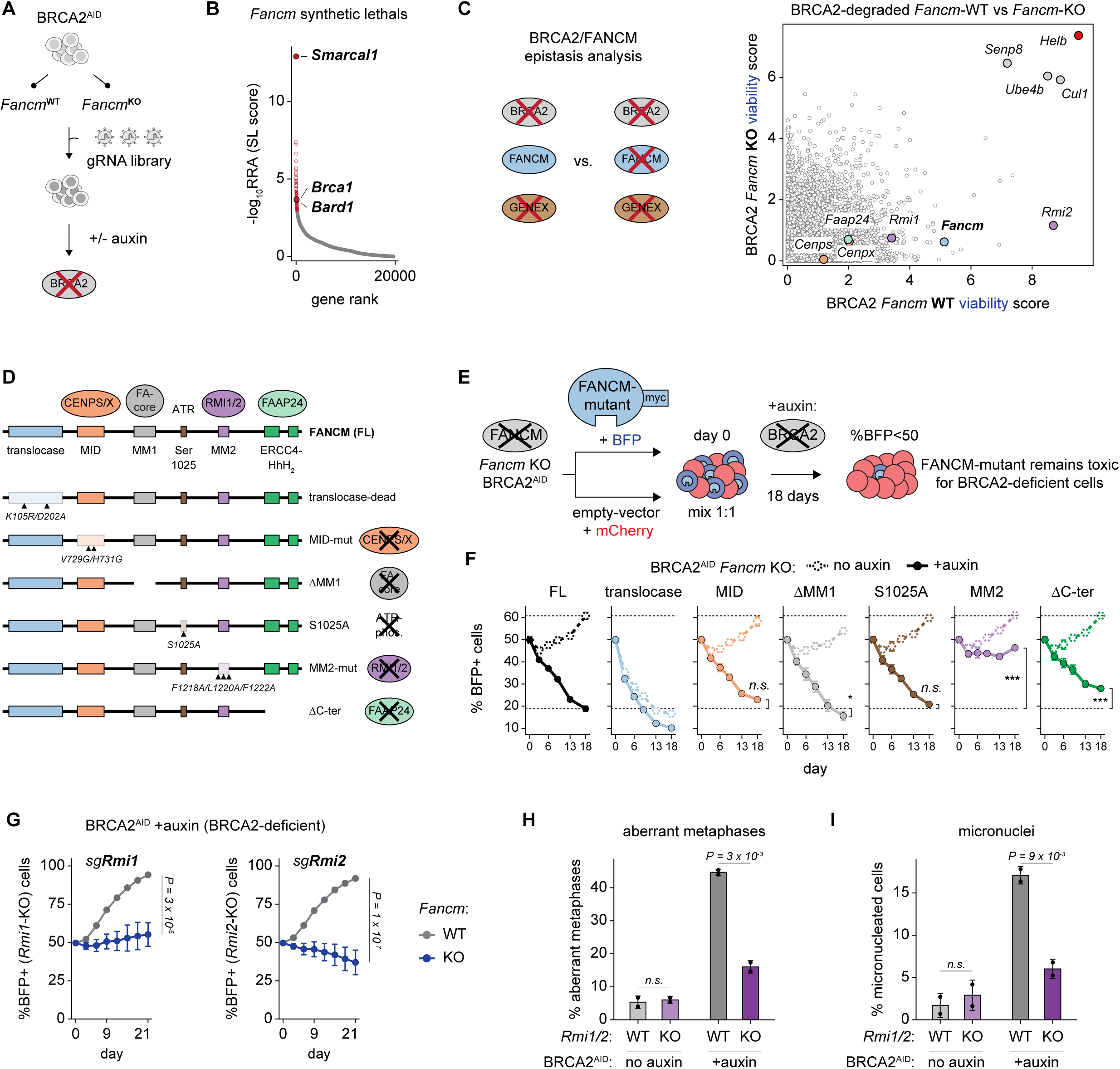
The FANCM-RMI1/2 complex promotes genome instability and toxicity in BRCA2-deficient cells. **A**, Schematic of the BRCA2-degron screen in *Fancm*-WT and *Fancm*-KO backgrounds. **B**, Dotplot of genes ranked according to their score of synthetic lethality with loss of *Fancm* (in BRCA2-proficient conditions). Significant synthetic lethal genes are shown as red circles. **C**, Scatterplot representing genome-wide synthetic viability interaction scores in BRCA2-depleted conditions, between *Fancm*-WT and *Fancm*-KO cells. **D**, Schematic of FANCM point and deletional mutants, and their impact on FANCM’s function. FANCM domains and physical interactors are shown as colored rectangles and circles, respectively. **E**, Schematic of the FANCM complementation fitness assay to identify FANCM’s toxic function. **F**, Competitive fitness assays during an 18-day time course of BRCA2-depletion (+auxin, thick line) in *Fancm*-KO clones, monitoring BFP-positive cells expressing the different FANCM-mutant transgenes shown in Fig. 3d. The mean ±SD of 2 independent clones is shown. Each FANCM-mutant is compared to FANCM-FL (brackets). Statistical analysis: Tukey’s HSD test; n.s. not significant; *P<0.05; ***P<0.001. **G**, Competitive fitness assays during a 21-day time course of BRCA2-depletion in *Fancm*-WT (grey) and *Fancm*-KO (blue) clones, monitoring BFP-positive cells expressing gRNA-pairs targeting *Rmi1* (left) or *Rmi2* (right). The mean ±SD of 2 independent clones is shown. Statistical analysis: two-way ANOVA. **H**, Barplots representing the fraction of cells with aberrant metaphases or **I**, micronuclei following 5 days of BRCA2-depletion in *Rmi1/2*-WT or *Rmi1/2*-KO cells. **H** and **I**, represent the mean ±SD of 2 independent clones. n≥100 metaphases or cells were scored. Statistical analysis: two-sided Student’s t-test.

We then investigated whether certain synthetic viable interactions with BRCA2-loss were shared or differed between the FANCM-proficient and the FANCM-deficient states (**Figure 3C**). *Helb* remained the top BRCA2-synthetic viable gene in both *Fancm*-WT and *Fancm*-KO cells (**Figure 3C**), indicating that the positive genetic interaction between *Helb* and *Brca2* (see **Figure 1** and **Figure S1**) is genetically independent of *Fancm*. Other shared synthetic viable hits were the cullin *Cul1*, the E3-ligase *Ube4b* and the cullin deneddylase *Senp8* (**Figure 3C**), all of which play vital roles in targeting proteins for proteasomal degradation ^27^ and whose loss likely abrogates auxin-mediated BRCA2 degradation.

Importantly, none of the FANCM physical interactors (CENPS, CENPX, RMI1, RMI2 and FAAP24) identified as synthetically viable with BRCA2-loss in *Fancm*-WT cells were found in *Fancm*-KO cells (**Figure 3C**). This demonstrates that these factors act epistatically with FANCM to limit the growth of BRCA2-deficient cells.

FANCM is a large protein that combines an ATP-dependent DNA translocase activity with multiple physically and functionally distinct protein interaction domains (**Figure 3D**) ^28^. These include MID which interacts with CENPS/X; MM1 with the FA-core complex; MM2 with RMI1/2 (two subunits of the BLM-TOP3A-RMI1/2 (BTR) complex); the ERCC4-like C-terminal domain with FAAP24; and a serine residue (S1025) that is phosphorylated by ATR (see **Figure 3D**). We sought to identify the FANCM domain most involved in mediating toxicity in BRCA2-deficient cells. Upon BRCA2-depletion in *Fancm*-KO cells, we reasoned that cells expressing a FANCM-mutant that retains its toxicity would be counter-selected, as opposed to a mutant that abolishes FANCM’s toxic function (**Figure 3E**). Thus, we complemented *Fancm*-KO cells with a series of point or deletional mutants of FANCM, each of which individually abrogates one of its aforementioned functions (**Figure 3D**) ^28^. Although FANCM-mutants were all expressed at similar levels (**Figure S5A**), the ΔMM1 (which disrupts interaction with the FA-core complex) and translocase-dead mutants failed to rescue the MMC-sensitivity of *Fancm*-KO cells (**Figure S5B, C**), as previously reported ^29–31^.

As expected, reintroducing full-length FANCM (FL) into *Fancm*-KO cells was toxic only in the context of BRCA2-depletion (**Figure 3F**). The ΔMM1 and S1025A mutants were also counter-selected, to the same extent as FANCM-FL (**Figure 3F**), indicating that interaction with the FA-complex and phosphorylation by ATR are not involved in FANCM’s toxic function. It should be noted that the translocase-dead mutant was equally toxic in BRCA2-proficient and -deficient cells (**Figure 3F**), consistent with the observation that inactivating mutations in FANCM’s translocase domain result in its toxic accumulation (“trapping”) on DNA ^30^. Strikingly, the MM2 mutant exhibited virtually no toxicity in BRCA2-deficient cells, whereas the MID and ΔC-ter mutants displayed slightly reduced toxicity compared to FANCM-FL. This is consistent with *Rmi1*/*Rmi2* being higher-scoring synthetic viable hits than *Faap24* and *Cenps/x* in the BRCA2-screen and validation experiments (**Figure 1** and **Figure S1**). Together, these results indicate that interaction with RMI1/2 through the MM2 domain is the primary cause of FANCM toxicity in BRCA2-deficient cells.

In competitive fitness assays, loss of *Rmi1*/*Rmi2* was only synthetically viable with BRCA2-loss in *Fancm*-WT cells, but not in Fancm-KO cells (**Figure 3G**). This confirms that RMI1/2 fully depend on FANCM to mediate toxicity in BRCA2-deficient cells. We further generated *Rmi1*-KO and *Rmi2*-KO clones in the BRCA2-degron background (**Figure S5D**), which displayed increased sensitivity to methyl methanesulfonate (**Figure S5E**), as expected ^32^. As with *Fancm*, knocking out *Rmi1* or *Rmi2* rescued genomic instability in BRCA2-deficient cells, as determined by decreased levels of aberrant metaphases and micronucleated cells (**Figure 3H, I**). Taken together, our results reveal that the FANCM-RMI1/2 complex promotes genome instability and toxicity in BRCA2-deficient cells, and that disrupting this axis is sufficient to restore normal cellular proliferation in the absence of BRCA2.

BRCA2 plays a major role in DSB repair by HR by loading the RAD51 recombinase. We therefore investigated whether loss of *Fancm* restores HR-repair and RAD51 foci formation in BRCA2-deficient cells. We used a modified version of the DR-GFP substrate ^33^, in which the repair of an I-SceI induced DSB restores functional BFP fluorescence only if repaired by HR (**Figure 4A**). This DR-BFP substrate was stably integrated in *Fancm*-WT and KO cells, prior to transfecting these cells with I-SceI (**Figure S6A**). Importantly, the I-SceI gene was linked to a mCherry cassette, so that only I-SceI expressing cells would be scored, thereby accounting for differences in transfection efficiency between samples that would artefactually bias measurements of HR-efficiency (**Figure S6A**). As expected, I-SceI transfection led to the appearance of BFP-fluorescent cells, indicative of efficient HR-repair of the I-SceI mediated DSB (**Figure 4B**). Degradation of BRCA2 resulted in a significant decrease in HR-efficiency, to similar levels between *Fancm*-WT and *Fancm*-KO cells (**Figure 4B**). Furthermore, RAD51-foci were virtually undetectable following BRCA2-depletion in both *Fancm*-WT and *Fancm*-KO cells (**Figure 4C, D** and **Figure S6B, C**). Together, these experiments demonstrate that inactivation of FANCM does not rescue HR-dependent DSB-repair in BRCA2-deficient cells. In addition to its role in HR, BRCA2 is also essential for protecting stalled replication forks from nucleolytic degradation, which leads to chromosome breaks and rearrangements in BRCA2-deficient cells ^34,35^. To monitor fork degradation, we sequentially labelled replicating DNA with CldU and IdU before treating cells with hydroxyurea (HU) to stall replication forks (**Figure 4E**). Depleting BRCA2 resulted in nascent strand degradation as assessed by the shortening of the IdU tract relative to the CldU tract (**Figure 4E, F**). Remarkably, knocking-out *Fancm* in BRCA2-deleted cells restored fork protection, to levels similar to those observed in BRCA2-proficient cells (**Figure 4E, F**). Similarly, fork protection was also restored in *Rmi1*- and *Rmi2*-KO BRCA2-deficient cells (**Figure 4G**). Collectively, these results demonstrate that loss of the FANCM-RMI1/2 axis rescues fork degradation, but not HR, to promote genome stability and proliferation in BRCA2-deficient cells.

**Figure 4:**
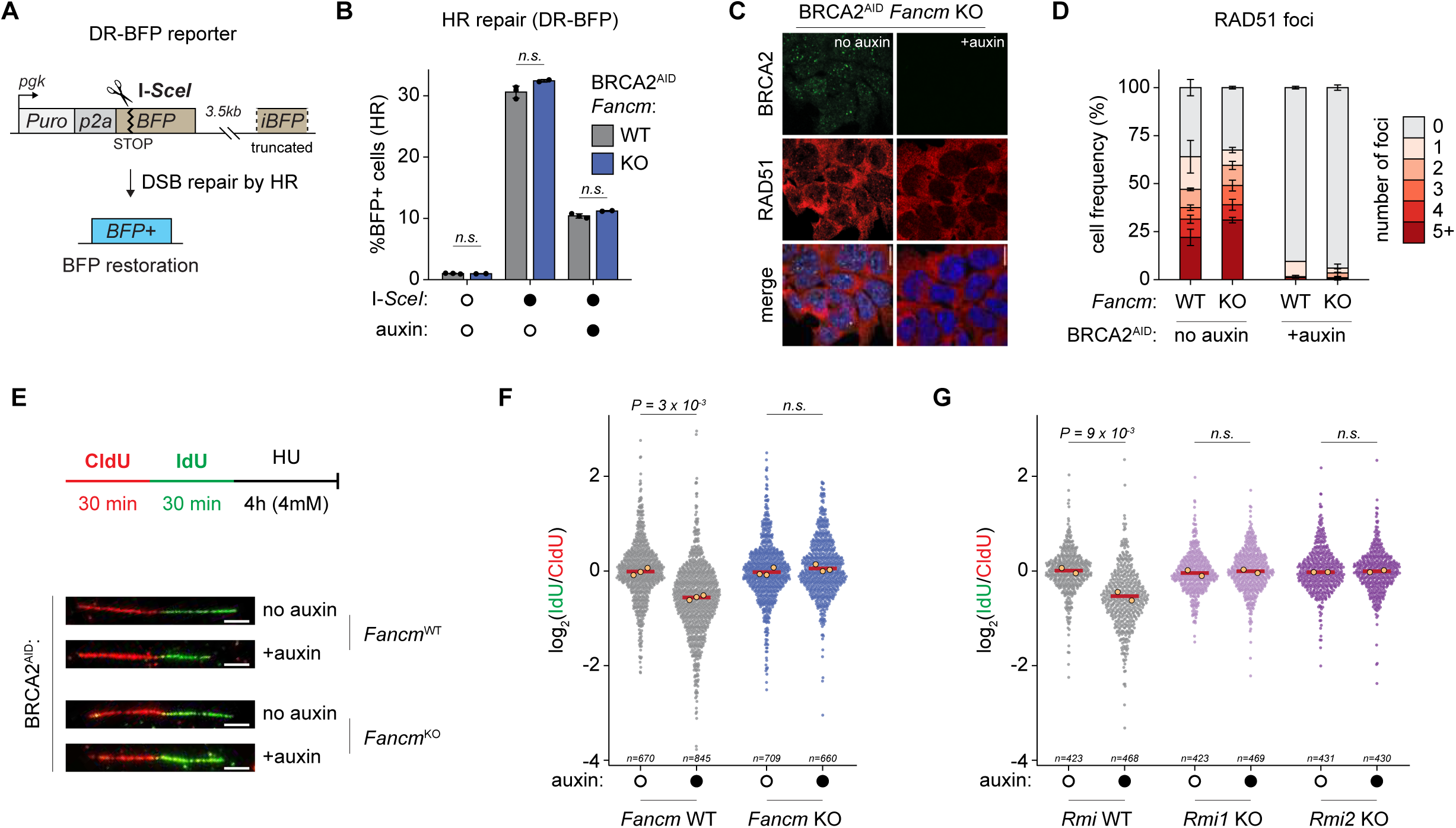
Loss of FANCM-RMI1/2 restores replication fork protection but not HR in BRCA2-deficient cells. **A**, Schematic of the DR-BFP reporter used to monitor HR-mediated repair of an I-SceI-induced DSB. **B**, Barplots representing the fraction of BFP-positive cells, corresponding to cells that have undergone HR-mediated repair, following Piggybac-mediated integration of the DR-BFP reporter and subsequent I-SceI expression in *Fancm*-WT and *Fancm*-KO cells, in contexts of BRCA2-proficiency (no auxin) or deficiency (+auxin). Data represent the mean ±SD of 2 independent clones. n≥2×10^5^ cells were scored by flow cytometry. Statistical analysis: two-sided Student’s t-test. **C**, Representative images of RAD51 foci in *Fancm*-KO cells subjected to 5 days of BRCA2-depletion. **D**, Barplots representing the fraction of *Fancm*-WT or *Fancm*-KO cells with 0, 1, 2, 3, 4 or ≥5 RAD51 foci after 5 days of BRCA2-depletion. Data represent the mean ±SD of 2 independent clones. n=100 cells were scored. **E**, Schematic of the labeling/treatment scheme to monitor replication fork degradation, and representative DNA fiber images of the ensuing replication tracts. **F**, Dotplot with quasirandom offsetting representing the IdU/CldU ratio of individual DNA fibers scored in WT, *Fancm*-KO and **G**, *Rmi1*-KO or *Rmi2*-KO cells following 2 days of BRCA2-depletion. **F** and **G**, the median IdU/CldU ratio of each individual experiment is shown as a larger yellow dot while the average of these medians is shown as a thick red bar. The number of fibers scored in each condition is indicated at the bottom of the graph. Statistical analysis: two-sided paired Student’s t-test.

Conversely, our screens and validation experiments highlight that FANCM is essential for the survival of BRCA1-deficient cells (**Figure 1** and **3**), which is consistent with previous reports of *Fancm*/*Brca1* synthetic lethality ^30,36^. To further explore the nature of this synthetic lethality, *Fancm*-KO clones were now generated in the BRCA1-degron background (**Figure 5A and Figure S7A, B**). Strikingly, BRCA1-depletion in *Fancm*-KO cells resulted in severe increases in aberrant metaphases, micronuclei and 53BP1 foci formation as compared to *Fancm*-WT cells (**Figure 5B-F, Figure S7C**), with a significant proportion of metaphases showing two or more chromosomal abnormalities, including fusion (*e.g.* radials, dicentric) and breakage events (**Figure S7D**). Together, these results demonstrate that aberrant DSB-repair underlies the *Fancm*/*Brca1* synthetic lethality and reveal an essential role for FANCM in maintaining genomic integrity of BRCA1-deficient cells. Similar to our prior observations (**Figure 2**), *Fancm*-loss in BRCA1-proficient settings had no noticeable impact on chromosomal aberrations nor micronuclei formation (**Figure 5C, E** and Figure **S7D**) but did result in a higher proportion of cells with 53BP1 foci (**Figure 5D, F**).

**Figure 5:**
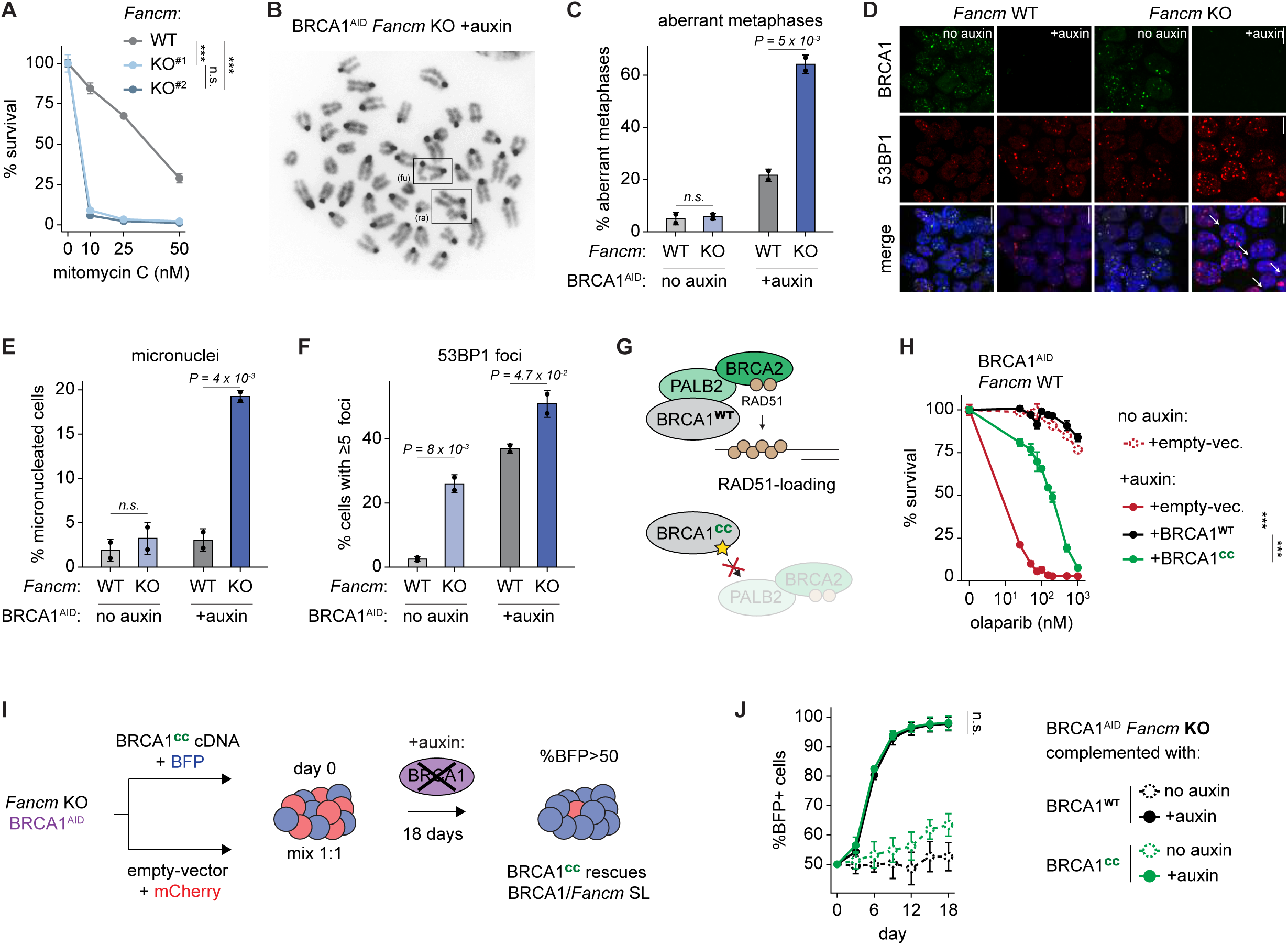
BRCA1 maintains genome stability in *Fancm*-deficient cells independently of PALB2/BRCA2. **A**, Cell survival after 5 days of treatment with increasing concentrations of MMC in *Fancm*-WT and two independent *Fancm*-KO clones, in the BRCA1-degron background. The mean ±SD of 2 replicates is shown. Statistical analysis: Tukey’s HSD test; n.s. not significant; ***P<0.001. **B**, Representative image of a metaphase spread following 5 days of BRCA1-depletion in *Fancm*-KO cells. Sister chromatid fusions (fu) and radial chromosomes (ra) are visible. **C**, Barplots representing the fraction of cells with at least one aberrant metaphase **D**, representative images of micronuclei (white arrows) and 53BP1 foci **E**, barplots representing the fraction of cells with micronuclei and **F**, more than five 53BP1 foci in *Fancm*-WT and *Fancm*-KO cells upon BRCA1-depletion for 5 days (auxin). Data in **C**, **E** and **F**, represent the mean ±SD of 2 independent clones. n=60 metaphases were scored for aberrations, while n≥100 cells were scored for micronuclei/53BP1 foci. Statistical analysis: two-sided Student’s t-test. **G**, Schematic of BRCA1-mediated recruitment of PALB2/BRCA2, and its abrogation in the BRCA1-CC (coil-coil) mutant. **H**, Cell survival following 5 days of BRCA1-degradation (auxin) and olaparib treatment in *Fancm*-WT cells complemented with empty-vector, BRCA1-WT or BRCA1-CC cDNA. The mean ±SD of 2 independent experiments is represented. Statistics as in Fig. 5a. **I**, Schematic of the competitive growth assay to investigate whether BRCA1-CC can rescue *Fancm/Brca1* synthetic lethality. **J**, Competitive fitness assays during an 18-day time course of BRCA1-depletion, monitoring BFP-positive *Fancm*-KO cells complemented with BRCA1-WT or BRCA1-CC transgenes. The mean ±SD of 2 independent clones is represented. Statistics as in Fig. 5a.

An important function of BRCA1 is to recruit BRCA2, through a bridging interaction with PALB2, which in turn loads the RAD51 recombinase and initiates HR (**Figure 5G**). Yet paradoxically, our work reveals that *Brca1* and *Brca2* are implicated in opposite, synthetic lethal and viable interactions with *Fancm*, respectively. We therefore asked whether *Brca1*/*Fancm* synthetic lethality involves BRCA1 functions that are independent of BRCA2. To this end, we used a BRCA1 separation-of-function mutant harboring a substitution in the coil-coil domain, BRCA1^L1407P^, hereafter referred to as BRCA1-CC, which retains all the functions of BRCA1 except for its ability to interact with PALB2 and thus recruit BRCA2 (**Figure 5G**) ^37^. BRCA1-degron cells were complemented with the BRCA1-CC transgene or its WT counterpart, in both *Fancm*-WT and *Fancm*-KO backgrounds. While BRCA1-CC and BRCA1-WT were expressed at similar levels (**Figure S7E**), BRCA1-CC failed to restore complete olaparib resistance upon endogenous BRCA1-depletion (**Figure 5H**), consistent with previous reports ^38^. Remarkably however, BRCA1-CC fully rescued the viability of BRCA1-depleted *Fancm*-KO cells, to the same extent as BRCA1-WT (**Figure 5I, J**). These results indicate that the synthetic lethality between *Brca1* and *Fancm* involves functions of BRCA1 that are genetically separable and independent of BRCA2, and thus provide a rationale for the unexpected opposite *Fancm*/*Brca1* and *Fancm*/*Brca2* genetic interactions uncovered in our screens.

In light of our results, we speculated that *FANCM* would be essential in human cancer cells that have adapted to BRCA1 deficiency but not those that have adapted to BRCA2 deficiency. To test this hypothesis, first, we interrogated the DepMap dataset of genetic dependencies across a large panel of human cancer cell lines (**Figure 6A**). Following stratification of *BRCA1*-mutated, *BRCA2*-mutated and *BRCA*-proficient cancer cells (**Figure 6A**), *FANCM* (and its interactor *FAAP24*) was indeed highly essential for the fitness of *BRCA1*-mutated cells specifically (**Figure 6B, C**). USP1 and WDR48 (see **Figure 1**) were also detected as essential only in *BRCA1*-mutated cancer cell lines (**Figure 6B, C**, see Discussion). Conversely, *POLQ* appeared as a synthetic lethal target in *BRCA2*-mutant but not *BRCA1*-mutant cells (**Figure 6B, C**), as predicted by our BRCA-degron screens (**Figure S2D**, see Discussion). Also consistent with our screens (**Figure S2D**) and previous reports ^39^, *CIP2A* was detected as highly essential in both *BRCA1*- and *BRCA2*-mutated cancer cells (**Figure 6B, C**).

**Figure 6:**
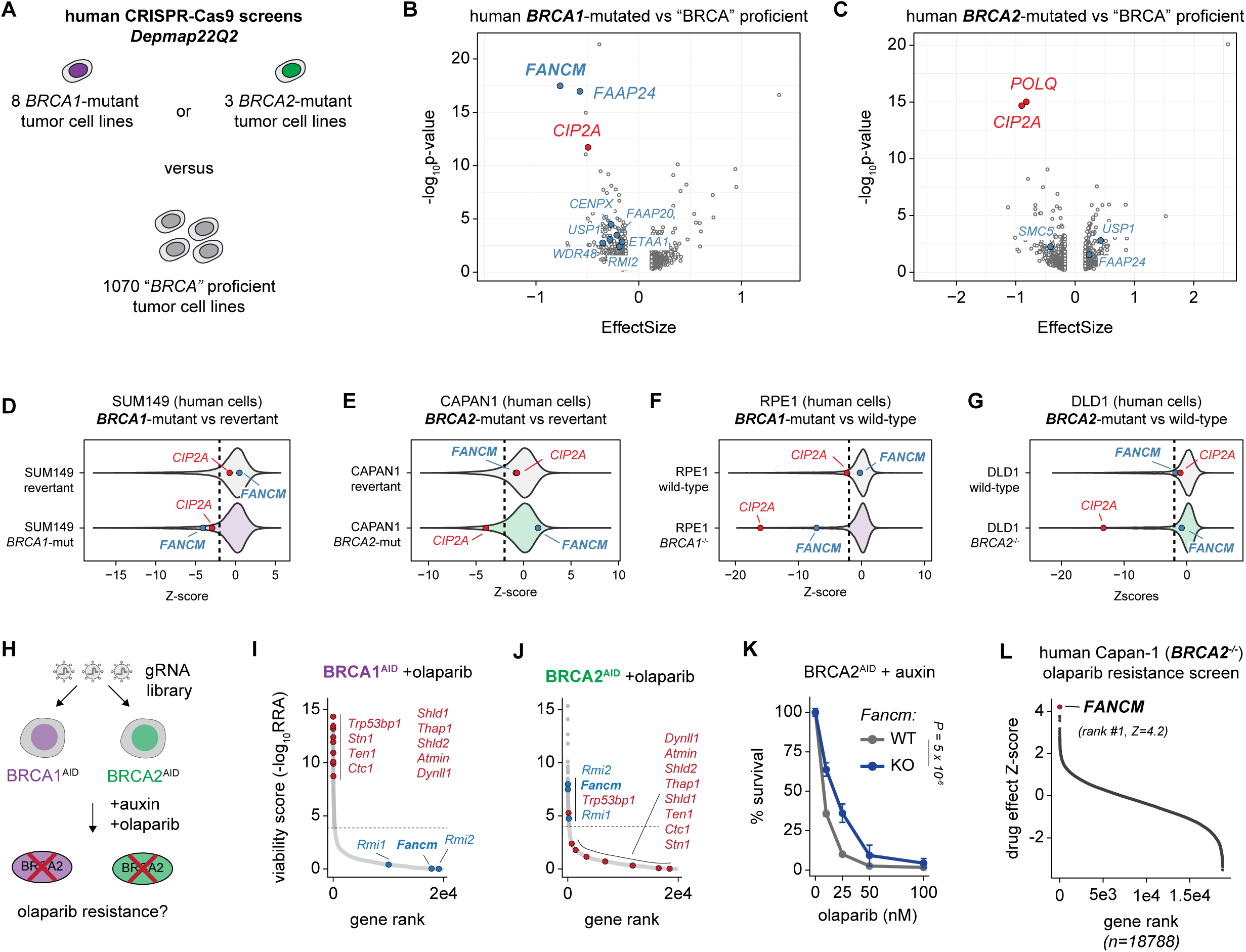
FANCM promotes PARPi sensitivity in *BRCA2*-deficient cells. **A**, Schematic of the analysis of DepMap genome-wide CRISPR-Cas9 screen data. Tumour cell lines from DepMap22Q2 release (PMID: 39468210) were classified as having either deleterious BRCA1 mutations (8 tumour cell lines), deleterious BRCA2 mutations (3 tumour cell lines) or no deleterious mutations in either BRCA1 or BRCA2 (1070 tumour cell lines). Using the class comparison tool (https://depmap.org/portal/interactive/custom_analysis) either BRCA1 or BRCA2 synthetic lethal genes were identified by comparing gene dependencies in either BRCA1 mutant or BRCA2 mutant tumour cell line cohorts to those in the cohort of tumour cell lines with no deleterious mutations in either BRCA1 or BRCA2. **B**, Volcano plots illustrating BRCA1 or **C**, BRCA2 synthetic lethal effects identified from DepMap analysis described in **A**. Negative effects (x-axis) indicate synthetic lethal effects, positive effects indicate synthetic viable effects. **D**-**G**. FANCM synthetic lethal effects identified in four genome-wide CRISPR-Cas9 screens. Violin plots of CRISPR-Cas9 screen data described in PMID: 40033056 are shown. Negative Z-scores represent synthetic lethal effects. CIP2A is shown as a positive control, being synthetic lethal in all four screens. FANCM is highlighted as a synthetic lethal in SUM149 and RPE *TP53*^−/−^ *BRCA1*^−/−^ screens (both *BRCA1*-mutant) but not in CAPAN1 nor DLD1 *BRCA2*^−/−^ screens (both *BRCA2*-mutant). **H**, Schematic of the olaparib resistance screen in BRCA1- and BRCA2-degron cells. **I**, Rank-ordered dotplot representing genome-wide viability scores (RRA) under olaparib treatment in BRCA1- or **J**, BRCA2-depleted cells. In **I** and **J**, the significance threshold (FDR<0.001 and RRA<0.001) is shown as a dotted line, genes involved in DNA-end protection are shown as red circles, while *Fancm/Rmi* are shown as blue circles. **K**, Cell survival after 5 days of BRCA2-depletion and treatment with increasing concentrations of olaparib in *Fancm*-WT (grey) and *Fancm*-KO (blue) cells. The mean ±SD of 2 independent clones is represented. Statistical analysis: two-way ANOVA. **L**, Rank ordered Drug Effect Z-score data from a CRISPR-interference screen for PARP inhibitor resistance in human *BRCA2*-mutant Capan1 cells. Capan1 cells were transduced with a genome-wide CRISPR interference sgRNA library and then exposed to 2000 nM olaparib for three weeks, a concentration that reduced the population size by 80 %, SF_20_ (see Methods). Gene-centric Drug Effect Z scores from 18,788 genes are shown, with sgRNA targeting *FANCM* causing the most profound PARPi resistance-causing effect.

We then reanalyzed a series of genome wide CRISPR screens ^39,40^ conducted in different *BRCA1*- and *BRCA2*-mutant human cell lines paired with their revertant or WT counterparts (**Figure 6D-G**). In all these screens, *CIP2A* was reproducibly identified as a synthetic lethal hit in BRCA-deficient cells, irrespective of *BRCA1*- versus *BRCA2*-mutation status. However, *FANCM* remained synthetically lethal only with BRCA1 loss, not BRCA2 loss. Notably, *FANCM* scored positively in the *BRCA2*-mutant Capan1 cancer cells, suggesting that loss of FANCM may benefit the growth of certain human BRCA2-deficient cancer cells.

Next, we carried out another series of genome-scale CRISPR screens in isogenic BRCA1- and BRCA2-degron cells exposed to a low dose of the clinically approved PARP inhibitor (PARPi), olaparib (**Figure 6H**). While DNA end-protection factors including members of the 53BP1-Shieldin-CST axis and DYNLL1/ATMIN were all identified as the principal drivers of PARPi-sensitivity in BRCA1-deficient cells (**Figure 6I**), these factors did not show significant effects in BRCA2-deficient cells (**Figure 6J**). Remarkably, *Fancm*, *Rmi1* and *Rmi2* were however identified as PARPi-sensitivity hits specifically in the BRCA2-degron screen (**Figure 6I, J**). Consistently, *Fancm*-KO cells exhibit decreased sensitivity to olaparib upon BRCA2-removal (**Figure 6K**) and were re-sensitized to PARPi upon FANCM reintroduction (**Figure S8A**). Finally, we performed a parallel CRISPR-interference screen for olaparib resistance in *BRCA2*-mutant human Capan-1 tumor cells, which identified FANCM as the most profound sensitivity factor (**Figure 6L**). Together, these results reveal that FANCM promotes PARPi sensitivity in BRCA2-deficient cells, while DNA-end protection factors act as key sensitizers of PARPi in BRCA1- but not BRCA2-deficient cells.

## DISCUSSION

Although BRCA1 and BRCA2 proteins are essential for viability due to their critical roles in HR and genome replication, genetically unstable cancer cells harboring *BRCA1* or *BRCA2* mutations can still sustain high proliferative capacity. To investigate this long-standing paradox, we established a genetic adaptation screening platform to identify genes that enable healthy cells to adapt to the loss of BRCA1 and BRCA2. Unlike previous screens carried out in cells adapted to BRCA1/2 deficiency, *i.e.* established tumor cell lines or clonally selected knockout cells, we performed parallel CRISPR-Cas9 screens in mouse embryonic stem cells engineered for acute (within hours) loss of endogenous BRCA1 or BRCA2, thereby tracing the adaptation routes that make initially healthy cells permissive to BRCA1 and BRCA2 deficiencies. The full isogenicity of the BRCA-degron cells enabled direct comparison of genetic interactions based on magnitude (score) and sign (negative versus positive) between BRCA1- and BRCA2-deficient cells, revealing hundreds of genes common to both backgrounds, specific to each, or acting in opposite ways.

In addition to its academic interest, our work is of clinical relevance as it may inform treatment strategies tailored to *BRCA1/2*-status. For instance, we find that *POLQ* exhibits a strong negative genetic interaction with *BRCA2* but not with *BRCA1*. Although POLQ has been shown to be essential for the survival of BRCA1- and BRCA2-deficient cells ^41,42^, our results are consistent with recent work showing that DNA end-resection is a critical determinant of POLQ-loss sensitivity in HR-deficient cells ^43^. Indeed, DNA end resection-defective *BRCA1*-null cells are less sensitive to POLQ-inhibition than *BRCA2*-null cells, in which resection is still active ^43^. Our genetic adaptation maps also identified genes that exhibit opposite interactions with BRCA1 and BRCA2. Notably, we found the USP1 deubiquitylase and its activator WDR48, which showed synthetic lethality with BRCA1 loss, are synthetic viable with BRCA2 loss. The essential role of USP1 in BRCA1-mutated cells has been previously reported ^44^, and USP1 inhibitors are under evaluation for clinical use in BRCA/HR-deficient cancers ^45,46^. Our findings indicate that inhibition of the USP1–WDR48 enzyme complex may be an effective therapeutic strategy for *BRCA1*-mutant, but not *BRCA2*-mutant, cancers ^47^.

Loss of resection antagonists is a well-established mechanism by which BRCA1-deficient cells regain viability and acquire resistance to PARPi. In line with this, 53BP1–SHLD–CST, DYNLL1/ATMIN, and HELB emerged as SV hits in the BRCA1-degron screen. Interestingly, aside from CST—which is essential for replication fork protection in BRCA2-deficient contexts ^14^—53BP1–SHLD, DYNLL1/ATMIN, and HELB were also identified as SV in BRCA2-deficient cells, with nevertheless noticeable score differences between BRCA1 and BRCA2. Indeed, in the BRCA1-degron screen, 53BP1–SHLD–CST and DYNLL1/ATMIN rank as top SV hits. Instead, in the BRCA2-degron screen, HELB is the top SV hit whereas 53BP1–SHLD and DYNLL1/ATMIN rank much lower. HELB also ranked first among SV genes in the BRCA2-degron screen conducted in the presence of olaparib, suggesting that loss of HELB may promote PARPi resistance in BRCA2-deficient cells. Competitive growth assays confirmed that the fitness benefit associated with 53BP1–SHLD loss is markedly greater in BRCA1-deficient cells than BRCA2-deficient cells. This disparity may reflect the distinct consequences of BRCA1 versus BRCA2 loss: in BRCA1-deficient cells, 53BP1–SHLD–CST/DYNLL1 promotes end protection and toxic NHEJ. Loss of these factors restores resection and permits generation of ssDNA substrates competent for RAD51 loading and HR ^15,16,48–55^. Conversely, in BRCA2-deficient cells, RAD51 filament formation is severely compromised despite active resection, effectively blocking HR progression. It therefore seems logical that loss of end protection via disruption of the 53BP1–SHLD/DYNLL1 axis confers only a mild benefit to BRCA2-deficient cells. HELB does not have end protection activity *per se* but does attenuates EXO1- and BLM-DNA2-dependent long-range resection via the binding of RPA-coated ssDNA ^56^. Once bound to the RPA-coated ssDNA complex, HELB can also catalyze RPA clearance from single-stranded DNA ^57^. The mechanism by which HELB loss benefits BRCA2-deficient cells remains unclear. One possibility is that increased long-range resection and accumulation of RPA-coated ssDNA in the absence of HELB create a permissive environment for the recruitment of alternative recombination factors that enable BRCA2-independent homology-directed repair.

We find that genes encoding for FANCM and its known physical interactors CENPS/MHF1, CENPX/MHF2, RMI1, RMI2, FAAP24, have opposing interactions with BRCA1 (*i.e.* SL) and BRCA2 (*i.e.* SV). FANCM has been reported to be essential for the survival of BRCA1-deficient cells ^30,36^, but we show here that this SL relationship implicates BRCA1 functions that are independent of PALB2/BRCA2. In contrast to the context of BRCA1 deficiency, we reveal that loss of FANCM improves the fitness of BRCA2-deficient cells at several levels. First, removal of FANCM restores normal proliferation in BRCA2-deficient cells. Second, loss of FANCM suppresses genetic instability (chromosomal aberrations and micronuclei formation) associated with BRCA2 deficiency. Third, Loss of FANCM decreases the sensitivity of BRCA2-deficient cells to the PARPi olaparib.

BRCA2 functions in HR and the protection of stalled replication forks are thought to contribute to genome integrity and chemotherapy response. We find that loss of FANCM does not restore RAD51 formation and HR in the absence of BRCA2. Instead, we show that loss of FANCM, or RMI1/2 (see below), significantly rescues fork protection in BRCA2-deficient cells upon fork stalling, to levels observed in wild type cells. Restoration of fork protection is a known driver of chemoresistance in BRCA2-deficient cells ^58^, and can be achieved by inactivating the fork reversing enzymes SMARCAL1, HLTF and ZRANB3 ^34,58,59^. However, these *bona-fide* fork-reversing enzymes were not identified in our BRCA2-degron screens, neither in steady state conditions nor in the presence of olaparib, suggesting that fork reversal might not be the main mechanisms through which FANCM promotes toxicity in BRCA2-deficient cells. In support to this, while FANCM can mediate fork reversal *in vitro* through its ATP-dependent translocase activity ^60^, protein domain-function analysis revealed that FANCM’s toxic activity in BRCA2-deficient cells is fully dependent on its MM2 domain (MM2), which mediates interaction with RMI1/2 ^31^. Epistasis between FANCM and RMI1/2 in BRCA2-deficient cells is remarkable, with the toxic activity of RMI1/2 only being observed in the presence of FANCM. RMI1/2 (through their interaction with BLM and TOP3A) and FANCM counter strand invasion by disrupting D-loops and dissolving recombination intermediates ^28,61,62^. Strand invasion is commonly used to rescue stalled replication forks ^63^. In the absence of BRCA2, FANCM–RMI1/2 may thus channel recombination intermediates into toxic rearrangements, further hindering the recovery of stalled replication forks and promoting genetic instability at the expense of cell fitness.

Overall, our data identify FANCM-RMI1/2 as a key determinant of cell fitness and PARPi response in BRCA2-deficient cells and uncover fundamental, previously underappreciated genetic and mechanistic differences in how cells adapt to the loss of BRCA1 versus BRCA2. These distinct adaptation pathways warrant further investigation to assess their potential in informing prevention and treatment strategies tailored to *BRCA1* or *BRCA2*-mutation status in clinical practice.

## ACKNOWLEDGEMENTS

We thank R.X. Coux, N. Festuccia and P. Navarro for sharing the E14tg2a mESC line and N. Johnson for sharing the BRCA1-L1407P cDNA. Work in L.D. laboratory was supported by Institut Pasteur, Institut Nationale de la Santé et de la Recherche Médicale and la Ligue Nationale Contre le Cancer (Labelisation 2019-2023 and 2024-2026) and Promex Stiftung für die Forschung. F.D. was supported by a Marie Skłodowska-Curie Actions postdoctoral fellowship (MSCA-PF# 101066732), a Pasteur-Roux-Cantarini postdoctoral fellowship and an EMBO Scientific Exchange Grant (SEG-11455).

## AUTHOR CONTRIBUTIONS

F.D. and L.D. conceived the study and experiments. F.D. performed the experiments unless stated otherwise, with the support of M.A. and F.M.Z. A.B. prepared the lentiviral and gDNA libraries under the supervision of R.M. F.F.S. conducted the Capan-1 screen under the supervision of C.J.L. F.D. conducted DNA fiber experiments under the supervision of D.G.A and M.L. C.J.L. and R.B. analyzed human cancer datasets. F.D. and L.D. wrote the manuscript, with input from all authors.

## METHODS

### Plasmid construction

The *TIGRE*-specific gRNA encoding plasmid (Addgene plasmid 92144) was a kind gift from Elphège Nora, while the hyperactive PiggyBac transposase (hyPBase) expression vector was a kind gift from Michel Wassef. All other plasmids originate from this study, and were assembled using restriction/ligation cloning.

The plasmid to simultaneously target CAS9 and OsTIR1(F74G variant) at the *TIGRE* locus (pFD155) was engineered by cloning *Cas9-t2a-hygro* and *OsTir1(F74G)-V5* cassettes under *pgk* and *CAG* promoters respectively, into a plasmid containing 5’ and 3’ homology arms for targeted insertion at *TIGRE* (Addgene plasmid 92141).

Targeting constructs to tag endogenous BRCA1 (pFD142) and BRCA2 (pFD143), respectively, with GFP-AID at their N-termini were generated as follows: ∼500bp homology arms (flanking both sides of, but excluding the start codons of *Brca1* and *Brca2*) were PCR-amplified from mouse genomic DNA. In the case of *Brca2*, at the PCR step, synonymous mutations were introduced within the 5’ end of the right homology arm, encompassing the seed and PAM sequences, so as to prevent CAS9-mediated cutting of the targeting vector upon transfection, and of the AID-tagged alleles following homologous recombination. Such mutagenesis was unnecessary for *Brca1* as the gRNA seed and PAM sequences are interspersed over both left and right homology arms. Left and right homology arms were then cloned, in frame, around a *blast^R^-GFP-AID* cassette, into a pUC19 backbone. Single gRNA expression plasmids (pFD144 and pFD146) to induce CAS9-dependent DSBs at the start codons of *Brca1* and *Brca2*, respectively, were cloned by ligating annealed DNA duplexes corresponding to the target gRNA sequences into BbsI-digested pX330 (gift from Feng Zhang, Addgene plasmid 42230).

The dual gRNA expression plasmids (used for fitness assays and generating KO clones) were cloned by assembling two sequential gRNA-expression cassettes, comprised of human and mouse U6 promoters, respectively, so as to avoid recombination-mediated excision of one of the two gRNA-expression cassettes. These sequential cassettes were cloned upstream of *pgk*-driven *puro-p2a-mCherry* (pFD165) or *puro-p2a-BFP* (pFD166) cassettes, within a vector containing PB transposon-specific inverted terminal repeats (gift from Michel Wassef). Importantly, BbsI-recognition sites yielding non-compatible overhangs were inserted immediately after both hU6 and mU6 promoters, enabling one-pot Golden Gate assembly of two different DNA duplexes (*i.e.* two different gRNA target sequences).

The BRCA1-CC (BRCA1^L1407P^) and BRCA1-WT expression constructs were generated by cloning human BRCA1^L1407P^ and BRCA1 cDNAs (gift from Neil Johnson) in frame with a C-terminal MYC-DDK tag, immediately downstream of a *CAG* promoter, within a vector containing a *pgk*-driven *puro-p2a-BFP* cassette and PB transposon-specific inverted terminal repeats. The myc-tagged FANCM-FL expression construct was cloned using an identical strategy (mouse FANCM cDNA from Origene, MR226436), into the same backbone as the BRCA1 constructs. FANCM-mutant constructs were then obtained by combining site-directed mutagenesis and/or overlap extension PCR, using myc-tagged FANCM-FL as a template. The empty vector (EV) used in both BRCA1 and FANCM cDNA expression experiments was generated by cloning a MYC-DDK coding sequence immediately downstream of a *CAG* promoter, within a vector containing a *pgk*-driven *puro-p2a-mCherry* cassette and PB transposon-specific inverted terminal repeats.

The DR-BFP plasmid (pFD181), used to monitor HR-repair of an I-SceI mediated DSB, was directly adapted from the pDRGFP reporter (gift from Maria Jasin, Addgene plasmid 26475). An I-SceI recognition site was inserted within the BFP ORF of a *pgk*-driven *puro-p2a-BFP* cassette, yielding premature stop codons and a truncated BFP. An “inactive” iBFP cassette corresponding to the ∼800bp surrounding the I-SceI site within the *puro-p2a-BFP* cassette was cloned 3.5kb downstream of the first cassette. This reporter system was then cloned between PB transposon-specific ITRs. The I-SceI expression vector (pFD182), enabling simultaneous expression of I-SceI and mCherry, was generated by inserting an *IRES-mCherry* cassette immediately after the I-SceI ORF of the pCBASceI vector (gift from Maria Jasin, Addgene plasmid 26477).

### Cell culture

All mESCs used in this study were derived from the E14TG2a mouse (strain 129/Ola) embryonic stem cell line (karyotype 40, XY; passage 19, kind gift from Rémi-Xavier Coux and Pablo Navarro). mESCs were grown on 0.1% gelatin-coated flasks or dishes, in 5% CO2 37°C incubators. For all experiments, cells were cultured in Serum + LIF conditions: DMEM+GlutaMax+Sodium pyruvate (Gibco), 15% ES-cell qualified FBS (Sigma), 0.1mM β-mercaptoethanol (Gibco), 1X non-essential amino acids (Gibco) and 1000 U/ml leukemia inhibitory factor (LIF, Chemicon). Unless stated otherwise, cells were passaged every other day using Trypsin (0.05%), and seeded at a density of 4-6×10^4^ cells/cm^2^. Cells were tested for mycoplasma contamination routinely (upon establishment of new clonal cell lines, and every month hereafter) and always tested negative. HEK293T cells were cultured in DMEM supplemented with 10% FBS, and 1% Penicillin/Streptomycin..

### Stable cell line generation

All genetically modified stable cell lines were generated using the 4D nucleofector X-unit (P3 primary cell reagents, CG-104 program, Lonza). CAS9-TIR1 expressing clones, and subsequent BRCA1^AID^ and BRCA2^AID^ cell lines were obtained by nucleofecting five million (parental) cells with 2.5ug of target gRNA-expressing plasmid and 15ug of non-linearized targeting donor (Maxipreps). Nucleofected cells were then serially diluted and plated on 10cm dishes. Two days later, antibiotic-dependent selection of integration events was initiated (hygromycin 300ug/mL, blasticidin 5ug/mL), and maintained for seven to ten days. Single colonies were then picked from plates showing ideal clonal density, and immediately split between one high-confluency plate used for PCR genotyping, and one low-confluency plate from which desired clones were further expanded until T25 density was reached, prior to being frozen as stable stocks. Stable knockout cell lines (*Fancm*-KO, *Rmi1*-KO, *Rmi2*-KO) were generated in a similar fashion, except that following nucleofection, gRNA-expressing cells were transiently selected (puromycin, 1ug/mL) for two days in T25 format, prior to being seeded (200 to 400 cells) in selection-free medium in a 10cm dish.

### Protein extraction and western blotting

Cells were trypsinized, resuspended in medium, washed once in PBS and immediately frozen at −80 °C as dry pellets. Pellets were then resuspended in ice-cold RIPA buffer (50 mM Tris-HCl pH 8.0–8.5, 150 mM NaCl, 1% Triton X-100, 0.5% sodium deoxycholate, 0.1% SDS) containing protease inhibitors (Roche), incubated for 30 min on ice and sonicated with Bioruptor (Diagenode) for three pulses of 15 seconds each. Lysates were then centrifuged for 20 min at 4 °C, and supernatants were kept. Protein concentration was determined using the Bradford (BioRad) assay. Samples were then boiled at 95 °C for 10 min in LDS buffer (Thermo) containing 200 mM DTT. For all western blots, 15-20ug of protein extracts were loaded for SDS-PAGE on 3–8% Tris-acetate polyacrylamide gels (Invitrogen), and transfer was performed on a 0.45-μm nitrocellulose membrane using a wet-transfer system, at 350 mA for 1.25h at 4 °C.

### Genotoxic drug sensitivity assay

1×10^4^ cells were seeded in 96-well format, in medium containing different concentrations of genotoxic drugs (Olaparib, MMC, MMS) in the presence of 5-Ph-IAA (Auxin analog, 1uM) to trigger BRCA1/2-depletion when necessary. Cells were counted 4-5 days later.

### Cumulative population doubling assays

3.5×10^4^ or 7×10^4^ cells were seeded in 48- or 24-well plate format respectively, counted and re-seeded at the same constant density every 2.5-3 days. Population doubling (PDL) was calculated at every passage as the log2 of the ratio between the number of cells counted, and the number of cells initially seeded. The cumulative population doubling was computed as the cumulative sum of all previous PDLs. Doubling time was then deduced by dividing the total elapsed time by the cumulative PDL.

### gRNA library cloning and lentiviral preparation

The mouse Brie CRISPR knockout lentiGuide-Puro pooled library was a gift from David Root and John Doench (Addgene #73633 ; http://n2t.net/addgene:73633 ; RRID:Addgene_73633), and was amplified using the protocol described by Sabatini et al. In the afternoon of D-1, 1.28×10^6^ HEK293T cells were seeded into two multilayer flasks (Falcon). Transfection was carried out the next day, after medium renewal. 69.7 µg of transfer plasmid, 55.9 µg of psPAX2 (Addgene #12260) and 20.2 µg of pMD2.G (Addgene #12259) were mixed with 7290 µL of 150 mM NaCl. Another mix of 875 µL of 1 mg/mL PEI MAX (Polysciences) and 7290 µL of 150 mM NaCl was made and both mixes were incubated at RT for 5min and then pooled together. The transfection mix was thoroughly vortexed and incubated at RT for 15 min before adding it to the cells. The supernatant was harvested at 48 and 72h post transfection filtered (0.45 µm), concentrated on Amicon ultra centrifugal filters (Merck), aliquoted, snap-frozen in liquid nitrogen and stored at -70°C.

### Genome-wide CRISPR screening in mESCs

Throughout the entire screen duration, Hygromycin (50ug/mL) was supplemented in culture medium so as to ensure continuous selection of CAS9-expressing cells. BRCA-degron mESCs were thawed in T25 format and amplified three days later by passaging 1.4×10^7^ cells into two T150 flasks. Three days later (i.e. one day prior lentiviral infection), 1.44×10^8^ cells were seeded in twenty 150mm dishes (Falcon).

The next day (Infection day, T0), cells were lentivirally transduced with the mouse genome-wide pooled gRNA knockout library (Brie, Addgene #7632) at multiplicities of infection (MOI) ranging between 0.25 and 0.45, in medium containing polybrene (8ug/mL, Sigma). The day after infection (T1), selection of transduced cells was initiated by replacing the old medium with fresh medium containing Puromycin (1ug/mL) in all but one plate, with selection-medium being refreshed daily thereafter. Infection efficiencies were determined two days later (T3) as the ratio of cell counts between one of the dishes under Puromycin selection and the “non-selected” dish. Multiplicities of infection (MOI) were then computed as follows: –ln(1 – infection_efficiency).

Four days post-transduction (T4), Puromycin-resistant cells were collected from the remaining eighteen dishes, and selection was prolonged for another three days by seeding 7×10^7^ of these cells in ten 150mm dishes containing Puromycin (1ug/mL). Seven days post-transduction, puromycin selection was stopped, and the screen was initiated as follows: For “control” conditions, 5.5×10^7^ cells were seeded in batches of seven 150mm dishes containing either DMSO (0.1%) or sublethal doses of Olaparib (LD_10_=50nM). For BRCA-depleted condition, the exact same “control” seeding and treatment conditions were used, except that the medium was supplemented with 5-Ph-IAA (Auxin analog, 1uM) to induce acute BRCA1 and BRCA2 degradation.

For each of these conditions, cells were passaged every three days (72 hours) in medium containing the necessary drugs, for a period of fifteen days in total, at a constant seeding density of 5.5×10^7^ in seven 150mm dishes. Upon each passaging days, aliquots of 1×10^8^ cells (or less in certain very selective conditions) were sampled and frozen for subsequent sequencing.

### gDNA library preparation and sequencing

Genomic DNA (gDNA) extraction was performed from dry cell pellets using QIAGEN Blood & Cell Culture DNA Maxi Kit following the manufacturer protocol. To maintain full-size library coverage, at least 80 million cells were used (coverage >500X). After the last ethanol wash, the DNA pellet was resuspended in 800µL of ultrapure H2O. Due to strong selection pressure in the context of olaparib treatment, some samples did not have 80 million cells available. All the cells recovered were used in that case. PCR amplification was performed using Herculase II (Agilent), by assembling multiple PCR-mixes (20 µL of 5X HerculaseII Buffer, 1µL of 10µM forward primer mix P5, 1 µL of 10 µM reverse indexed primer P7, 1 µL of 100 mM dNTP, 2 µL of Herculase II, H2O qsp 100µL) with the following cycling conditions: 95°C, 1 min; then 28 cycles of (95°C, 30s; 53°C, 30s; 72°C,30s) then 72°C, 10 min. For the full-size library samples, 28 reaction were conducted, using 10 µg gDNA in each case. For gRNA library sequencing, 10ng of plasmid library in one 100-µL reaction was used. PCR purification was performed with AMPure magnetic beads (Beckman). PCR replicates were pooled back together and 100 µL was collected for purification. 55 µL of beads were added and incubated for 15 min at RT (0.55X). Samples were placed on a magnet and amplicon-containing supernatant was transferred to a new tube. 45 µL of beads were added and incubated for 15 min at RT (1X). Once placed on the magnet, primer-containing supernatant was removed and beads washed twice with 200µL of 80% EtOH. EtOH was removed tubes were air dried for 1 min. Beads were resuspended in 16 µL ultrapure H2O and incubated for 5 min at RT. Tubes were placed back on the magnet to elute and remove beads from sample. Sample purity and concentration were assessed with DNA 5000 ScreenTape (Agilent). Another round of AMPure PCR purification was performed for samples sowing extra peaks. An equimolar pool was made based on TapeStation concentrations. Next-Generation-Sequencing (50bp single read) was performed on a NextSeq 2000 device (Illumina) using 10% PhiX (Illumina). Sequencing results and counting tables were processed using a tool developed by the bio-informatic core facility of Institut Curie ^64^.

### Two-color fitness assays

1×10^5^ cells were seeded in a 24-well plate, prior to being lipofected (LTX, Thermo) the next day with 300ng of HyPBase plasmid, and 900ng of dual gRNA expression plasmid. One day later, selection of transposition events (puromycin, 1ug/mL) was initiated and maintained for seven days, at which stage virtually all cells show blue (BFP, target gRNAs plasmid) or red (mCherry, control gRNAs plasmid) fluorescence. For each target gene tested, blue and red cells were mixed in 1:1 proportions, and 7×10^4^ cells of this mix were seeded in a 24-well plate, in the presence of Auxin and Olaparib when necessary. Cells were then passaged every three days, and the BFP/mCherry proportion was tracked by flow cytometry throughout timecourse duration. A similar strategy was used for fitness assays involving BRCA1-CC/WT and FANCM-mutant complemented cells.

### RNA extraction, reverse transcription and qPCR

RNA extraction was performed using the RNeasy kit and on-column DNase digestion (Qiagen). Reverse transcription was performed on 1 μg total RNA using SuperScript III (Invitrogen), in the presence of RNasin (Promega). To quantify transcript expression, qPCR was performed on diluted cDNA (1/80) using SYBR Green mastermix (Applied Biosystems). Relative mRNA expression was determined using the ΔΔCt method. The arithmetic mean of Ct values from two housekeeping genes, *Actb* and *Gapdh*, was used as the housekeeping Ct.

### Metaphase Analysis

Cells were cultured for 5 days (in the presence of auxin when necessary) and treated with 100ng/uL colcemid (KaryoMax, Gibco) for three hours prior to cell collection. Cells were then washed with PBS, subjected to hypotonic swelling in 75mM KCl at 37°C for 20 minutes and fixed in ice-cold methanol/acetic acid (3:1 v/v) fixative solution. Fixed cells in suspension were then dropped onto a humidified microscope slide from a height of 30 centimeters to facilitate the spreading of metaphase chromosomes. Slides were air-dried overnight and mounted in DAPI-containing mounting medium (ProLong Gold, Thermo). Metaphases were imaged using an Axio Imager Z2 microscope (Zeiss) and the automated imaging platform Metafer (MetaSystems).

### Immunofluorescence and micronuclei staining

Cells were cultured for 5 days (in the presence of auxin when necessary), and grown on Laminin (5ug/mL, Sigma) and Poly-L-Ornithine (15ug/mL, Sigma) coated coverslips for 2 days before being processed for Immunofluorescence. Cells were fixed on coverslips in PFA (4% in PBS) for 10 minutes at room-temperature, permeabilized in ice-cold permeabilization buffer (0.5% v/v Triton X-100 in PBS) for 5 minutes and blocked in blocking solution (1% w/v BSA in PBS) for 1 hour at room-temperature. Coverslips were then sequentially exposed to primary and secondary antibodies, diluted in blocking solution, for 1 hour at room temperature prior to being mounted in DAPI-containing mounting medium (ProLong Gold, Thermo). Images were acquired using an ECLIPSE Ti2 microscope (Nikon) and analyzed using ImageJ.

### Cell cycle analysis by EdU labeling and Propidium Iodine staining

2×10^5^ cells were seeded in 12-well format and labeled two days later with 10uM EdU for 45 minutes. Labeled cells were then trypsinized, washed once in PBS prior to being fixed in 4% PFA for 30 minutes at room temperature and washed once more in PBS. Click reaction between EdU and A647-azide (Invotrogen) was performed for 1h at room temperature, in the dark, by resuspending fixed cells in PBS containing 1mM CuSO4, 100mM L-ascorbic acid and 2uM A647-azide. Cells were then washed once in PBS and stained with Propidium Iodine (PI) overnight at 37°C by resuspending cells in PBS containing 0.5mM Tris-HCl, 0.75mM NaCl, 1.9mM Sodium Citrate, 250ug/mL RNAse A and 25ug/mL PI. Cells were then washed once in PBS and processed for flow cytometry.

### CRISPRi screen method

A positive selection, genome-wide CRISPR interference screen in Capan1 (*BRCA2*-mutant) cells was carried out in a dCas9-KRAB+ Capan1 derivative (generated using Lenti-dCas9-KRAB-blast, Addgene #89567), using the Weissman hCRISPRi-v2 sgRNA library (gift from Jonathan Weissman, Addgene #83969). The screen was carried out as previously described ^65^ with the exception that after transduction with the hCRISPRi-v2 library, cells were exposed to either olaparib (2000 nM, a concentration that reduced the population size by 80 %, SF_20_) or drug vehicle, DMSO, for three weeks. Sequencing of sgRNA from genomic DNA isolated from cells prior to and after olaparib exposure was used to estimate enrichment of sgRNAs. Pre-processing and quantification of sgRNA frequency was performed as previously detailed ^66^ to generate gene-centric Drug Effect Z scores (effect of CRISPRi of each gene on olaparib resistance) and also Viability Z scores (effect of of CRISPRi of each gene on cell survival in the absence of drug).

### HR-monitoring using DR-BFP

1×10^5^ cells were seeded in a 24-well plate, prior to being lipofected (LTX, Thermo) the next day with 300ng of HyPBase plasmid, and 1.5ug of DR-BFP reporter plasmid (pFD181). One day later, selection of transposition events (puromycin, 1ug/mL) was initiated and maintained for five days. Selected cells were then seeded in a 24-well plate (1×10^5^ cells), in the presence of auxin when necessary, to ensure BRCA2-depletion prior to inducing I-SceI expression. The next day, cells were lipofected with 1ug of I-SceI-IRES-mCherry plasmid (pFD182). BFP and mCherry fluorescence were assessed 72h later using flow cytometry.

### DNA fiber spreading

2×10^5^ cells were seeded in a 12-well plate, in the presence of auxin when necessary. 2 days later, cells were sequentially labeled with 90uM CldU and 225uM IdU for 30 minutes each, prior to being exposed to HU (4mM) for 4h. Cells were then trypsinized, counted and resuspended in PBS at a concentration of 7.5×10^5^ cells/mL. 2uL of cell suspension were mixed with 7uL of lysis buffer (200 mM Tris-HCl, pH 7.4, 50 mM EDTA, and 0.5% v/v SDS) on a microscope slide, and lysis was left to proceed for 5 minutes. Slides were then slowly tilted at a 15-30° to spread DNA fibers, prior to being air-dried and fixed for 2 minutes in ice-cold methanol/acetic acid (3:1 v/v) fixative solution. Fixed slides were then denatured for 1h in 2.5M HCl, washed thoroughly with PBS and blocked for 1h in blocking solution (0.1% Triton (v/v) and 1% BSA (w/v) in PBS). Slides were incubated for 2h in blocking solution supplemented with anti-BrdU primary antibodies recognizing CldU (1:200) and IdU (1:100), washed with PBS and lastly labeled for 1h in blocking solution supplemented with fluorescent secondary antibodies. Fibers were imaged with an ApoTome.2 microscope (Zeiss) and analyzed using ImageJ.

## SUPPLEMENTARY FIGURES

**Supplementary Figure 1: (for Figure 1).**
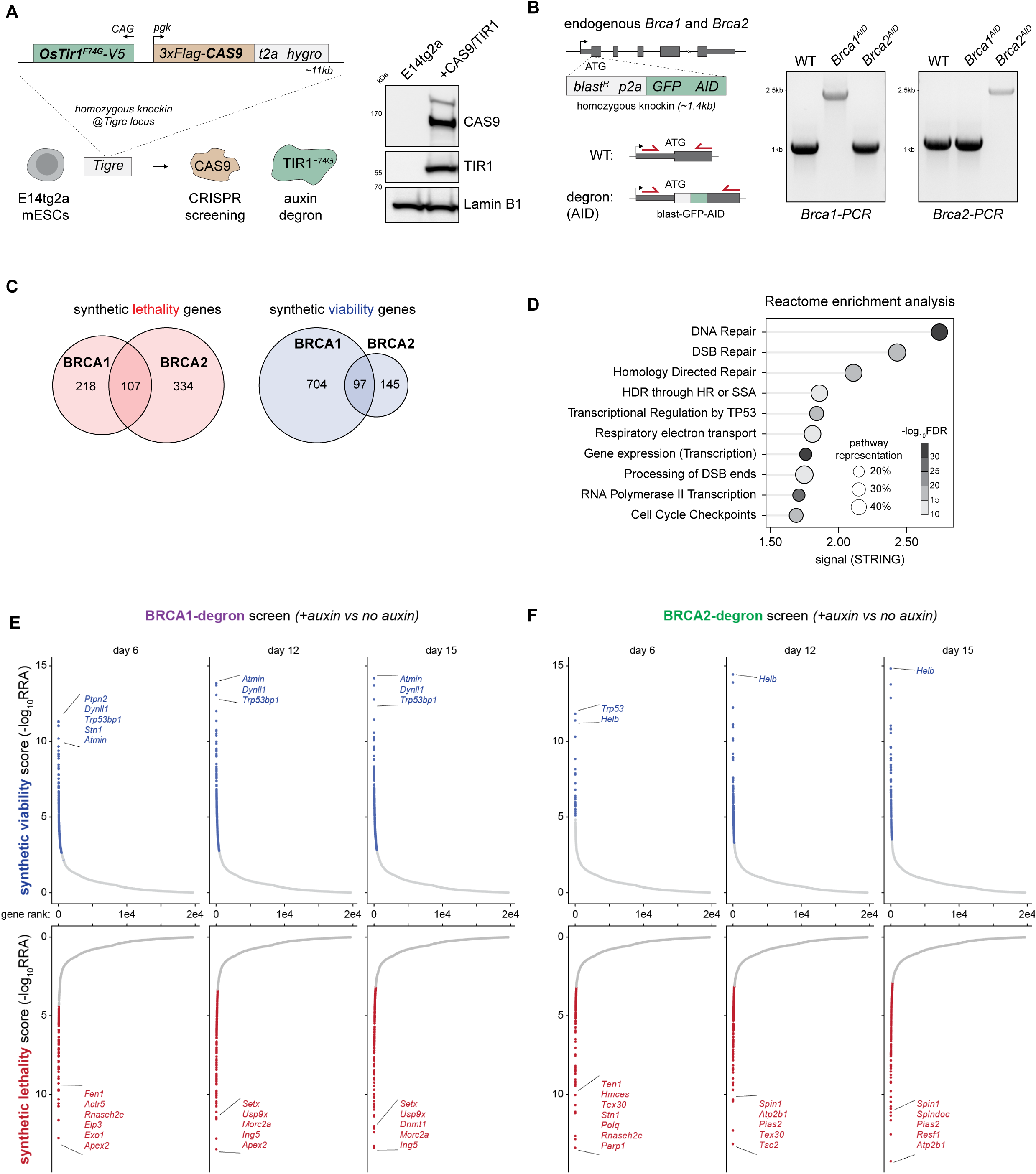
**a**, Targeted homozygous insertion of *CAS9* and *OsTIR1^F74G^* at the *TIGRE* locus (left) results in constitutive CAS9 and TIR1 protein expression as assessed by western blot (right). **b**, Schematic representation of homozygous knockin of GFP-AID at (and in frame with) the start codon (N-terminus) of endogenous *Brca1* or *Brca2*, and PCR-genotyping of the resulting BRCA1- and BRCA2-degron cell lines. **c**, Venn diagram representation of total synthetic lethal and viable interactions scored in the BRCA1- and BRCA2-screens (FDR<0.05 and RRA<0.05). **d**, Top 10 Reactome pathways enriched among total BRCA1- and BRCA2- screen hits, ranked by signal (FDR-adjusted p-values with Benjamini-Hochberg correction). **e**, Dotplot representation of genes targeted by the genome-wide gRNA library ranked according to their synthetic viability (top) or synthetic lethality (bottom) scores across three different timepoints in the BRCA1-degron and **f**, BRCA2-degron screens. Statistically significant synthetic viable and lethal genes are shown as blue and red dots, respectively. A few top genes are labeled.

**Supplementary Figure 2: (for Figure 1).**
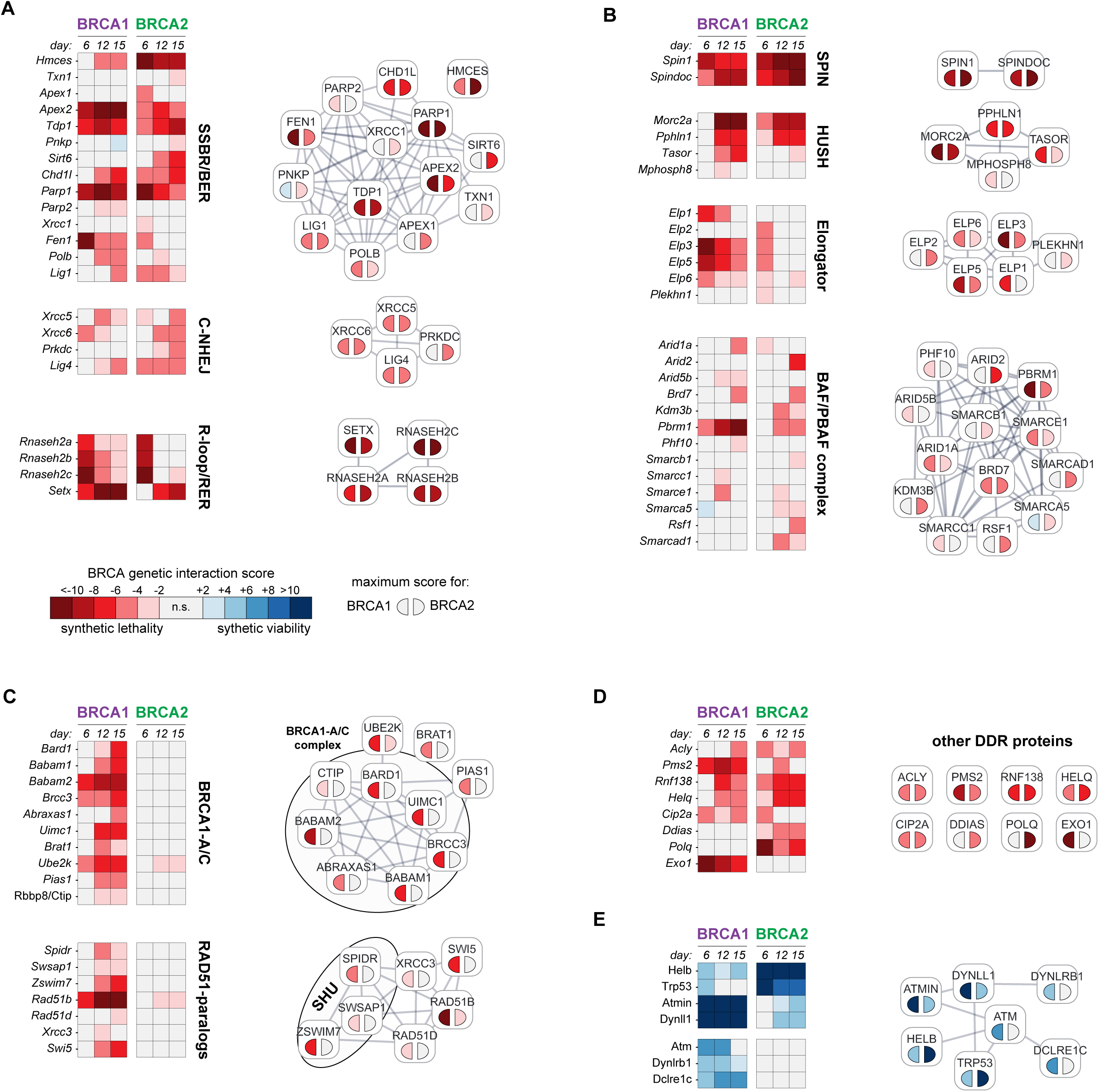
**a**, **b**, **c**, **d** and **e**, Heatmaps showing synthetic lethal (red shades) and viable (blue shades) interaction scores (MAGeCK RRA) for different genes at the three timepoints of the BRCA1-(left heatmap) and BRCA2- (right heatmap) screens. STRING interaction networks for each gene groups are shown on the right, with genes colored according to their maximal BRCA-interaction score in each screen.

**Supplementary Figure 3: (for Figure 1).**
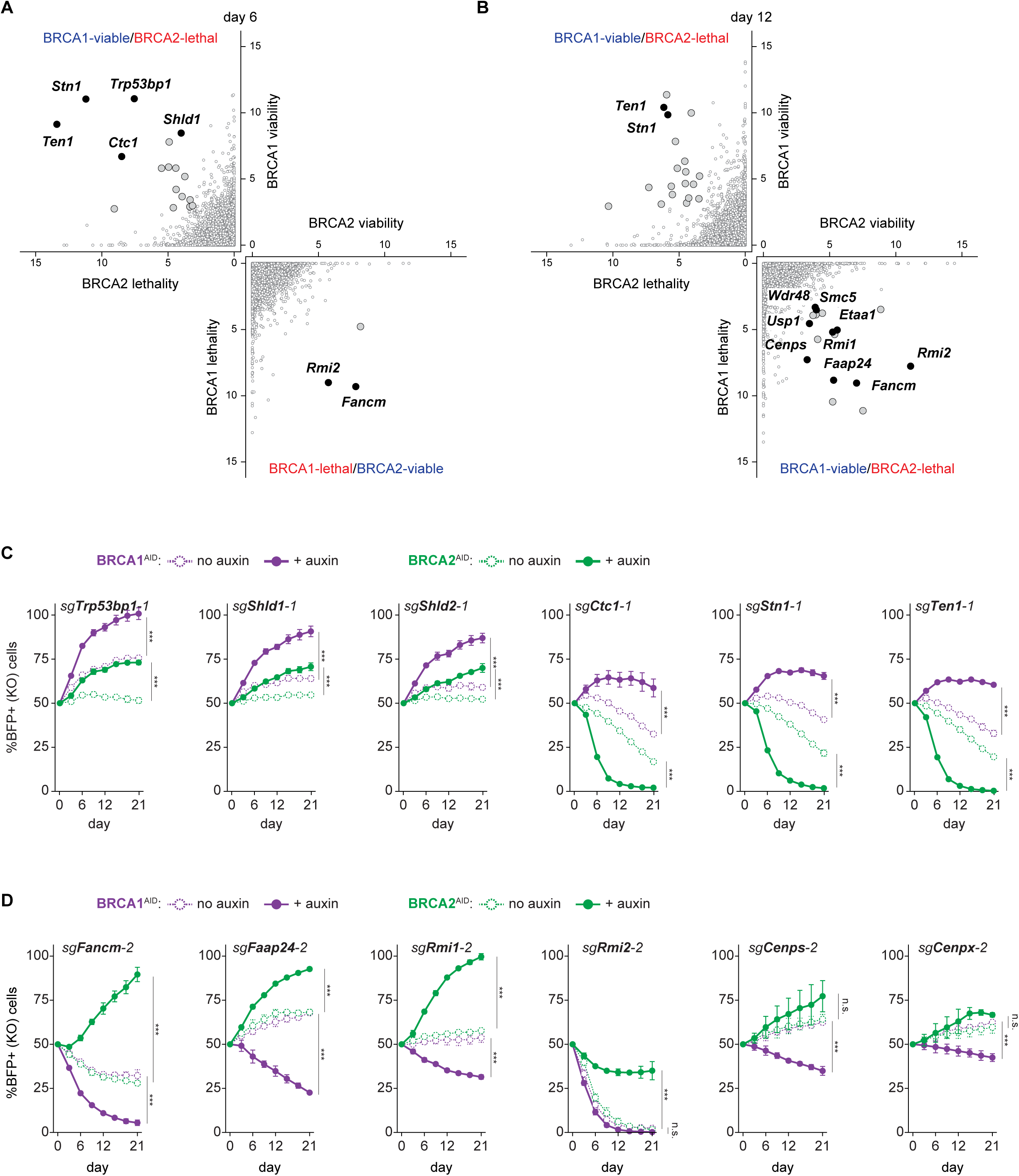
**a**, Scatterplot showing BRCA1-SV versus BRCA2-SL scores (top-left quadrant) and BRCA1-SL versus BRCA2-SV scores (bottom-right quadrant) at day 6 and **b**, day 12. Genes meeting significance threshold (FDR<0.05 and RRA<0.05) are represented as larger dots. **c**, Competitive fitness assays during a 21-day time course of BRCA1- (purple) or BRCA2-(green) depletion, monitoring BFP-positive cells expressing gRNA-pairs targeting 53BP1-Shieldin-CST components or **d**, FANCM and its physical interactors. **c** and **d**, represent the mean ±SD of 2 independent experiments. Statistical analysis: two-way ANOVA; ***P<0.001.

**Supplementary Figure 4: (for Figure 2).**
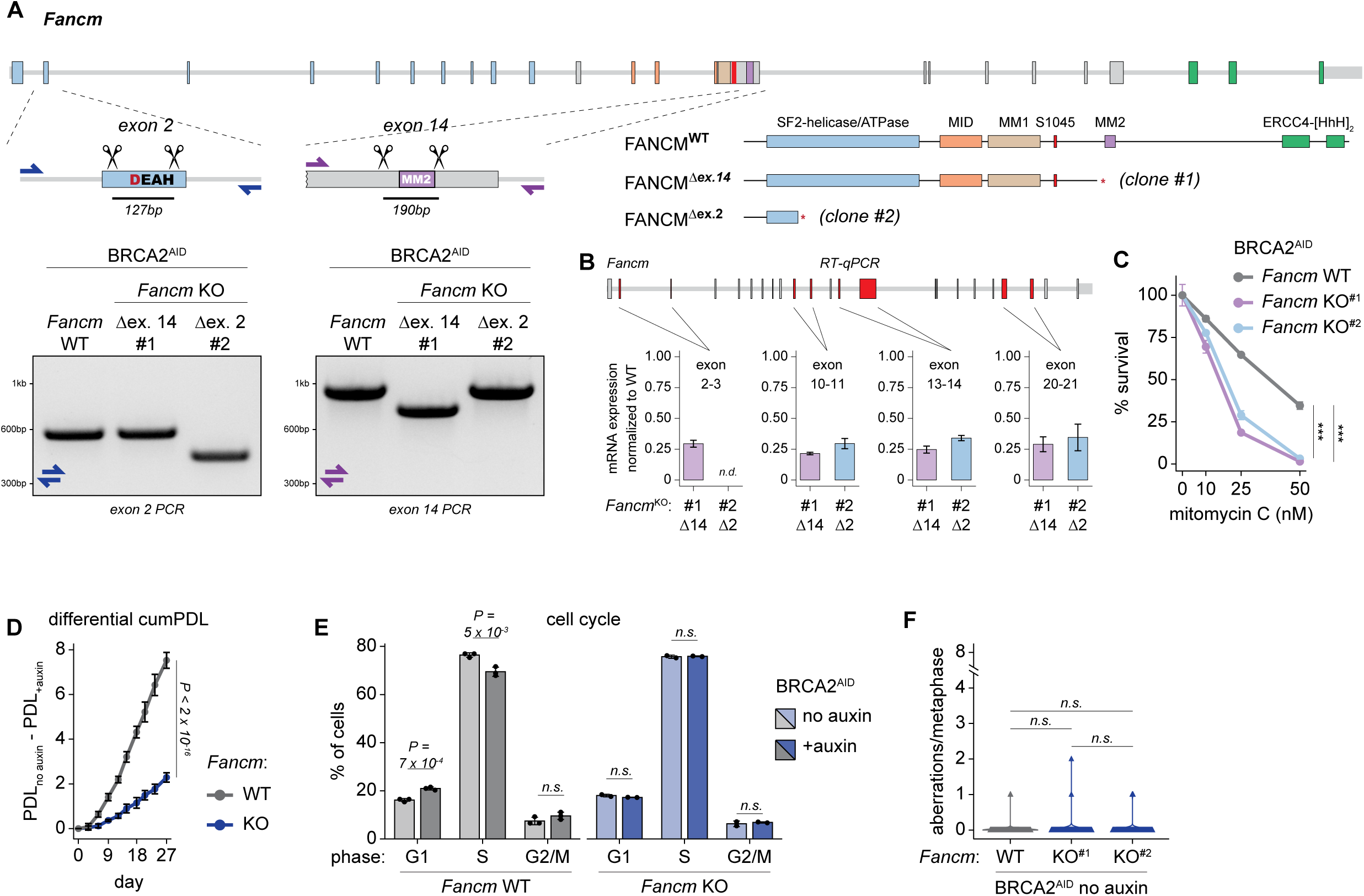
**a**, Schematic of the *Fancm* locus, with exons represented as larger rectangles, colored according to the different FANCM-domains they encode. gRNAs used to delete exon 2 or exon 14 of *Fancm* are represented as scissors, and PCR-genotyping of two independent *Fancm*-KO clones reveals homozygous deletion. The resulting FANCM-truncated products are also shown. **b**, qPCR analysis of *Fancm* mRNA expression reveals severely reduced transcript levels in both *Fancm*-KO clones, across 4 different exon-boundaries. mRNA expression was quantified with the ΔΔCt method, using two housekeeping genes (*Actin* and *Gapdh*) as reference genes, and *Fancm*-WT cells as reference sample. The mean ±SD of 2 independent replicates is represented. **c**, Cell survival after 5 days of treatment with increasing concentrations of MMC in *Fancm*-WT and two independent *Fancm*-KO clones, in the BRCA2-degron background. The mean ±SD of 2 replicates is shown. Statistical analysis: Tukey’s HSD test; n.s. not significant; ***P<0.001. **d**, Cumulative population doubling differences between BRCA2-proficient (no auxin) and BRCA2-depleted (+auxin) cells, during 27 day time course, in *Fancm*-WT (grey) and *Fancm*-KO (blue) clones. The mean ±SD of 2 independent clones is represented. Statistical analysis: two-way ANOVA. **e**, Cell cycle analysis in *Fancm*-WT and *Fancm*-KO cells, in BRCA2-proficient and BRCA2-depleted conditions. The mean ±SD of 2 or 3 independent experiments is represented. Statistical analysis: two-sided Student’s t-test. **f**, Violin plots representing the number of chromosomal aberrations per cell among metaphases scored in Fig. 2d, in BRCA2-proficient cells. The mean ±SEM of 150 metaphases is shown as a circle. Statistical analysis: two-sided Wilcoxon rank-sum test.

**Supplementary Figure 5: (for Figure 3).**
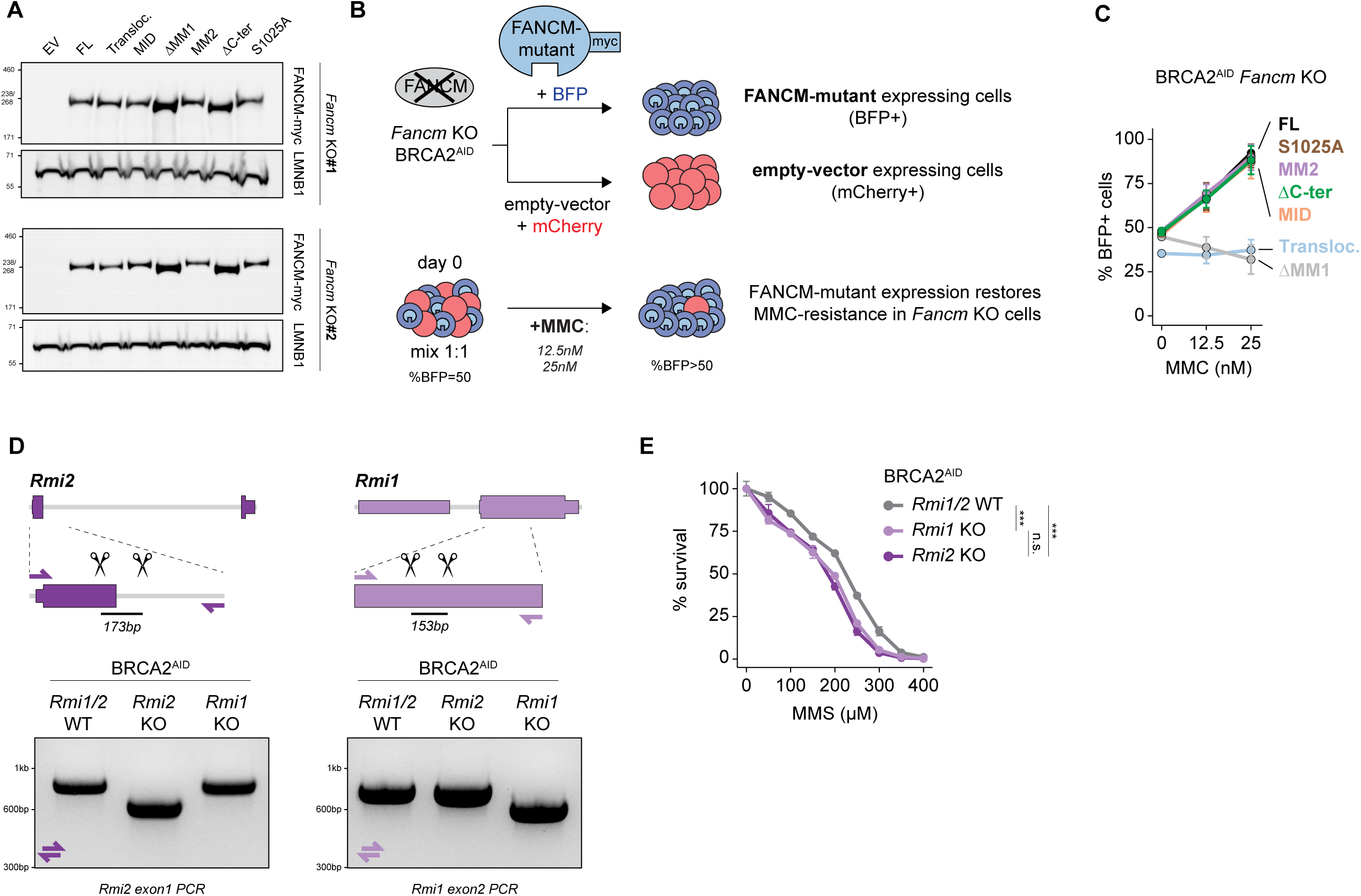
**a**, Western blot analysis of two independent *Fancm*-KO clones following Piggybac mediated integration of myc-tagged empty-vector (EV) or different FANCM-mutant transgenes. Membranes were probed with antibodies against myc and Lamin B1. **b**, Schematic of the FANCM complementation fitness assay to identify which FANCM-mutants rescue MMC sensitivity. **c**, Competitive fitness assays in *Fancm*-KO clones after 6 days of treatment with increasing concentrations of MMC, monitoring BFP-positive cells expressing the different FANCM-mutant transgenes shown in Fig. 3d. The mean ±SD of 2 independent clones is represented. **d**, Schematic of the *Rmi2* and *Rmi1* loci, with exons represented as larger rectangles. gRNAs used to delete *Rmi2* exon 1 or *Rmi1* exon 2 are represented as scissors, and PCR-genotyping of *Rmi2*-KO and *Rmi1*-KO clones reveals homozygous deletion. **e**, Cell survival after 5 days of treatment with increasing concentrations of methyl methanesulfonate (MMS) in *Rmi1/2*-WT, *Rmi1*-KO and *Rmi2*-KO clones, in the BRCA2-degron background. The mean ±SD of 2 replicates is shown. Statistical analysis: Tukey’s HSD test; n.s. not significant; ***P<0.001.

**Supplementary Figure 6: (for Figure 4).**
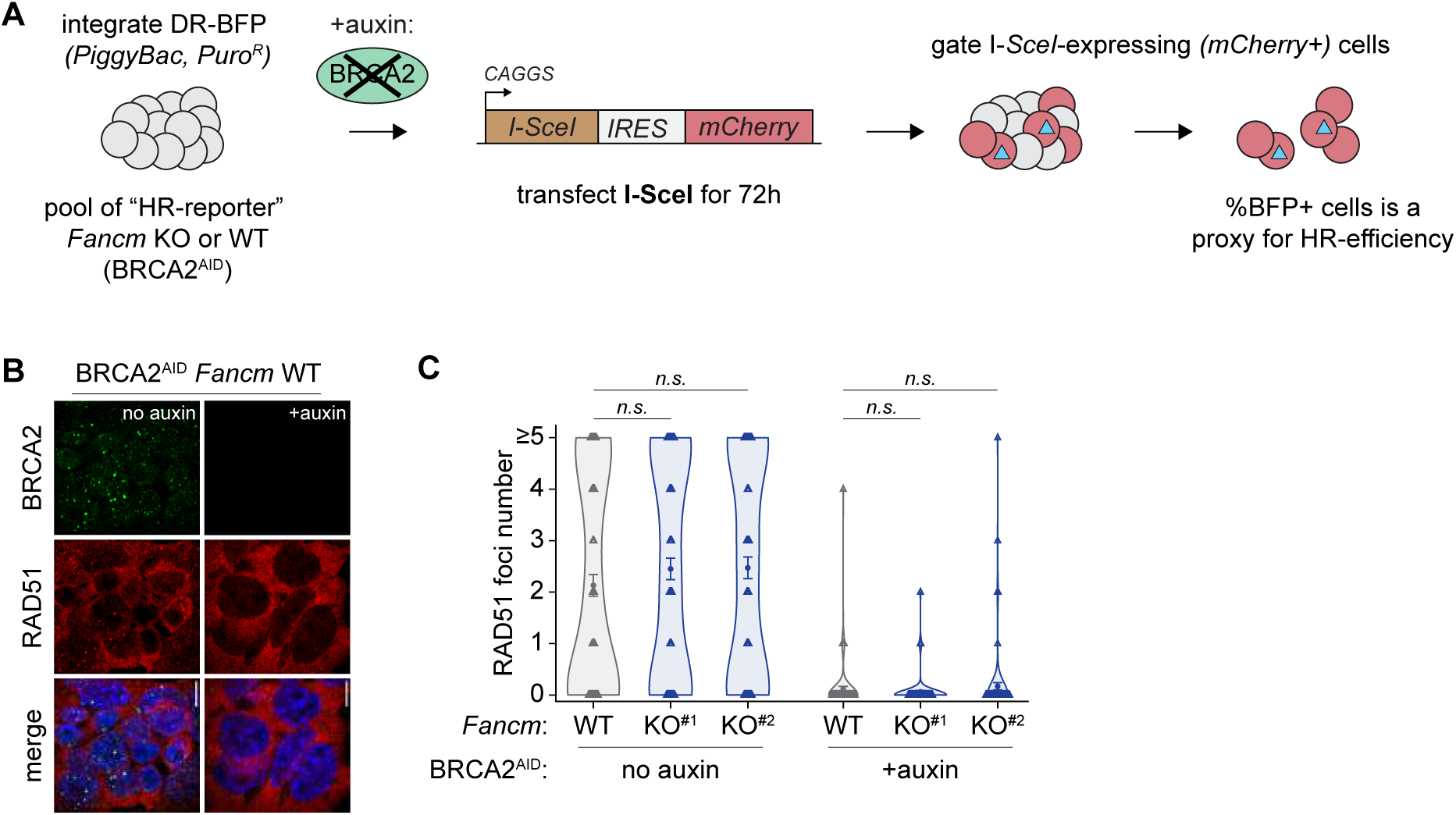
**a**, Schematic of the I-SceI expression and flow cytometry gating strategy to monitor HR-repair strictly in I-SceI expressing cells. **b**, Representative images of RAD51 foci in *Fancm*-WT cells subjected to 5 days of BRCA2-depletion. **c**, Violin plots representing the number of RAD51 foci among cells scored in Fig. 4d. The mean ±SEM of 100 cells is shown as a circle. Statistical analysis: two-sided Wilcoxon rank-sum test.

**Supplementary Figure 7: (for Figure 5).**
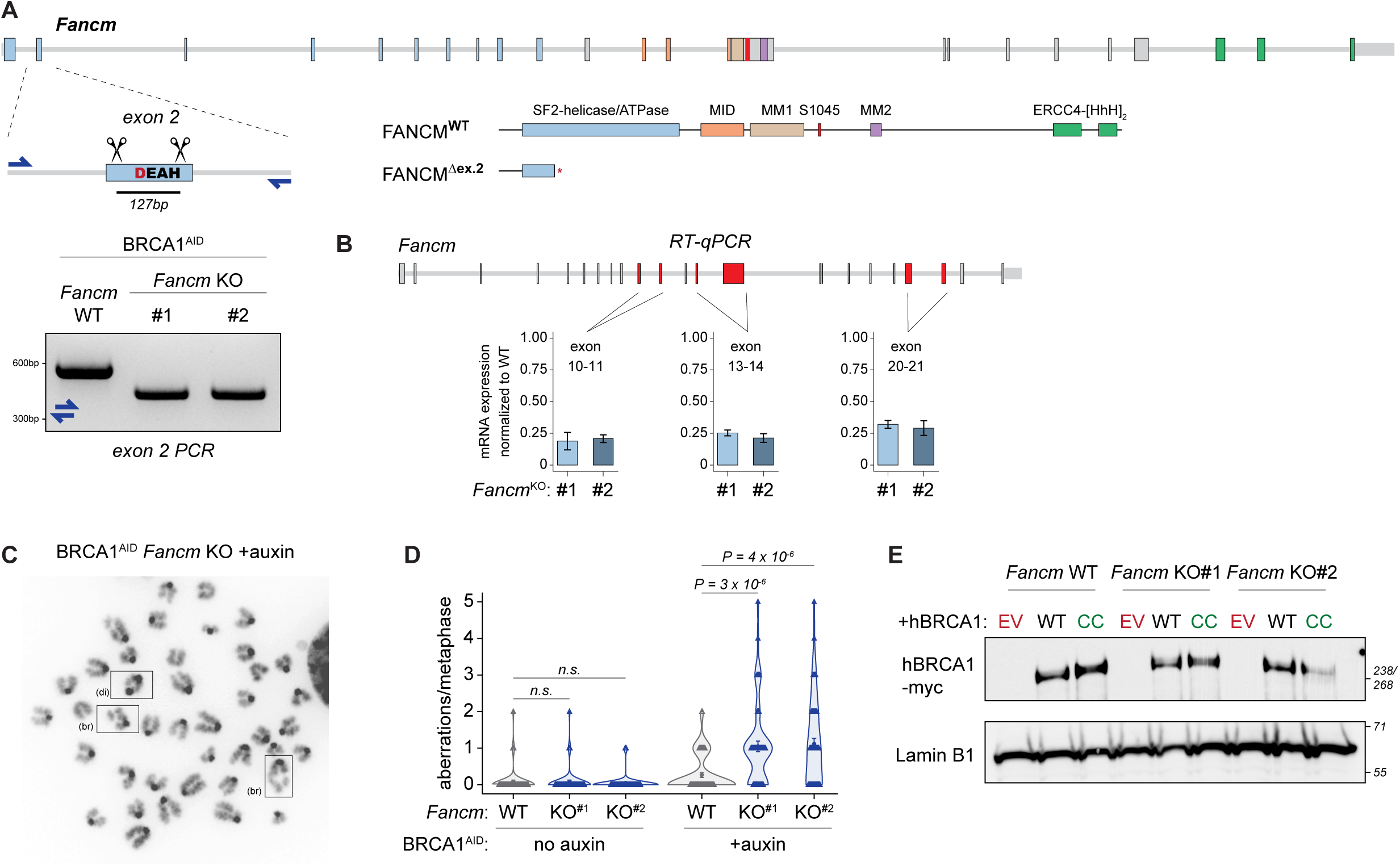
**a**, Schematic of the *Fancm* locus, with exons represented as larger rectangles, colored according to the different FANCM-domains they encode. gRNAs used to delete the exon 2 of *Fancm* are represented as scissors, and PCR-genotyping of two independent *Fancm*-KO clones reveals homozygous deletion. The resulting FANCM-truncated product is also shown. **b**, qPCR analysis of *Fancm* mRNA expression reveals severely reduced transcript levels in both *Fancm*-KO clones, across 3 different exon-boundaries. mRNA expression was quantified with the ΔΔCt method, using two housekeeping genes (*Actin* and *Gapdh*) as reference genes, and *Fancm*-WT cells as reference sample. The mean ±SD of 2 independent replicates is represented. **c**, Representative image of a metaphase spread following 5 days of BRCA1-depletion in *Fancm*-KO cells. Dicentric chromosomes (di) and chromatid breaks (br) are visible. **d**, Violin plots representing the number of chromosomal aberrations per cell among metaphases scored in Fig. 5c. The mean ±SEM of 60 metaphases is shown as a circle. Statistical analysis: two-sided Wilcoxon rank-sum test. **e**, Western blot analysis of *Fancm*-WT and two independent *Fancm*-KO clones following Piggybac mediated integration of myc-tagged empty-vector (EV), BRCA1-WT or BRCA1-CC transgenes. Membranes were probed with antibodies against myc and Lamin B1.

**Supplementary Figure 8: (for Figure 6).**
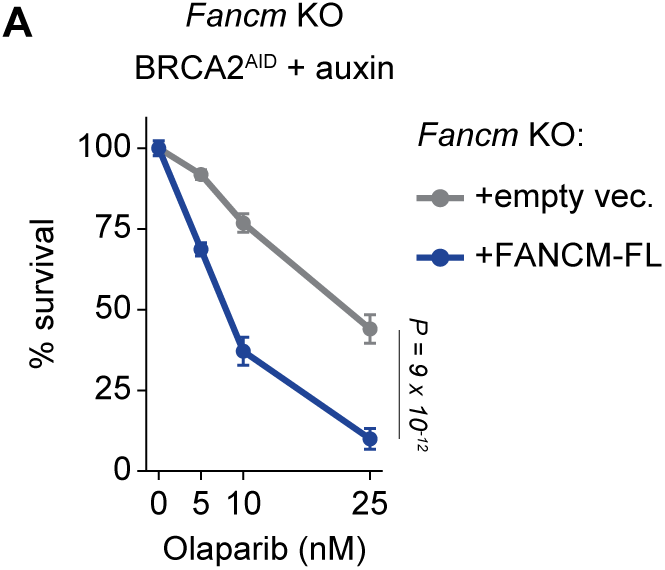
**A**, Cell survival after 5 days of BRCA2-depletion and treatment with increasing concentrations of olaparib in *Fancm*-KO cells complemented with empty-vector (grey) or FANCM-FL (blue) transgenes. The mean ±SD of 2 independent clones is represented. Statistical analysis: two-way ANOVA.

## SUPPLEMENTARY TABLES

**Supplementary Table 1: BRCA1- and BRCA2-degron screen analysis and scores.**

This table contains MAGeCK analysis results obtained after comparing DMSO (BRCA-proficient) vs Auxin (BRCA-deficient) treatment conditions. Each sheet is specific to one screen (BRCA1-degron or BRCA2-degron), one timepoint (D6, D12 or D15), one type of analysis (depletion/synthetic_lethality or enrichment/synthetic_viability) and are named accordingly. Targeted gene names, RRA scores, FDR, rank and log2FC are included, and are preceded by “neg” or “pos” terms, depending on whether a depletion (synthetic lethality) or enrichment (synthetic viability) analysis was conducted, respectively.

**Supplementary Table 2: Reactome enrichment analysis.**

Reactome enrichment analysis conducted on all significant hits identified in either of the BRCA1 or BRCA2 screen, at any of the three timepoints.

**Supplementary Table 3: List of genes involved in opposite genetic interactions with BRCA1 versus BRCA2.**

This table lists the name of genes involved in opposite genetic interactions with BRCA1 versus BRCA2, comprising 42 BRCA1-viable/BRCA2-lethal and 20 BRCA1-lethal/BRCA2-viable genes.

**Supplementary Table 4: List of primers.**

List of all oligonucleotides used in genotyping and RT-qPCR experiments.

**Supplementary Table 5: List of gRNAs.**

List of all gRNAs used to generate knock-ins and knock-outs.

**Supplementary Table 6: List of plasmids.**

List of all plasmids used in this study.

**Supplementary Table 7: List of antibodies.**

List of antibodies used in this study.

**Supplementary Table 8: Statistics summary.**

List of all statistical tests conducted in this study, with references to each figure panel where they are employed. For each test, assumptions are highlighted where relevant, and both the value of the statistic and its associated P-value are indicated.

## Supplementary text

The intrinsic isogenicity of the BRCA1/2-degron cells (**Figure 1A, E**) allows for direct comparison of the BRCA1- and BRCA2-centered genetic interaction networks, revealing that only a small fraction of synthetic lethal and synthetic viable genes are common to both networks (**Fig S1C**). Many of the shared BRCA1/2-synthetic lethal genes are involved in single strand break (SSB) detection, processing and repair during base excision repair (BER, **Figure S2A**), including *Apex2*, *Parp1*, *Tdp1*, *Hmces* and *Fen1*, which have previously been shown to be essential for the survival of BRCA1/2-deficient cancer cells ^1–6^. Consistent with previous studies ^7,8^, we recovered CIP2A and the three subunits of RNase H2 as synthetic lethal with both BRCA1 and BRCA2 (**Figure S2A, D**). Other notable shared BRCA1/2 synthetic lethals are the non-homologous end-joining (NHEJ) factors KU (*Xrcc5/6*), DNA-PKcs (*Prkdc*), and LIG4 (**Figure S2A**), suggesting an essential role for NHEJ in repairing DSBs throughout the cell cycle in HR-compromised mES cells. Interestingly, chromatin regulatory complexes including SPIN1/SPINDOC, HUSH, BAF/PBAF and most subunits of the tRNA-modifying ELP (elongator) complex were also identified in both screens (**Figure S2B**). These hitherto unreported synthetic lethal interactions could constitute therapeutic targets in BRCA1/2-mutated cancers.

We also uncovered several genes whose loss is synthetic lethal with BRCA1 but not with BRCA2, including subunits of the BRCA1-A/C complexes as well as RAD51-paralogs such as the SHU-complex (**Figure S2C**). The lack of synthetic lethality between BRCA2 and RAD51-paralogs is in agreement with epistasis between these factors ^9,10^. EXO1, which was recently shown to be essential for the survival of BRCA1- but not BRCA2-deficient cells ^11^, was accordingly identified as a BRCA1-specific synthetic lethal hit in our screen (**Figure S2D**). Conversely, POLQ exhibited a strong synthetic lethal interaction with BRCA2 but not with BRCA1 (**Figure S2D**, see discussion).

Screening upon acute loss of BRCA1/2 also enabled the concurrent detection of synthetic viable interactions, involving genes whose loss improves the fitness of non-adapted BRCA-deficient cells (**Figure S1E, F**). *Trp53*, whose knockout delays lethality in *Brca1-* and *Brca2*-null mouse embryos ^12^ and is frequently inactivated in BRCA1/2-mutated tumors ^13^, was identified as synthetic viable in both the BRCA1 and BRCA2 screens (**Figure S2E** and **Figure S1E, F**). DYNLL1, its transcriptional activator ATMIN, and HELB were identified as shared synthetic viable genes in both screens (**Figure S2E**). DYNLL1 and HELB are resection inhibitors whose losses restore HR and promote resistance to PARPi in BRCA1-deficient cells ^14,15^. Our screens suggest that these anti-resection factors could also mediate sensitivity to DNA-damaging chemotherapy in BRCA2-deficient cancer, but that their action may be differentially important depending on BRCA-status (see discussion).

## REFERENCES

1. Hakem, R. et al. The Tumor Suppressor Gene Brca1 Is Required for Embryonic Cellular Proliferation in the Mouse. Cell 85, 1009–1023 (1996).

2. Sharan, S. K. et al. Embryonic lethality and radiation hypersensitivity mediated by Rad51 in mice lacking Brca2. Nature 386, 804–810 (1997).

3. Findlay, G. M. et al. Accurate classification of BRCA1 variants with saturation genome editing. Nature 562, 217–222 (2018).

4. Khalizieva, A., Moser, S. C., Bouwman, P. & Jonkers, J. BRCA1 and BRCA2: from cancer susceptibility to synthetic lethality. Genes Dev. 39, 86–108 (2025).

5. Chen, C.-C., Feng, W., Lim, P. X., Kass, E. M. & Jasin, M. Homology-Directed Repair and the Role of BRCA1, BRCA2, and Related Proteins in Genome Integrity and Cancer. Annu. Rev. Cancer Biol. 2, 313–336 (2018).

6. Scully, R., Glodzik, D., Menghi, F., Liu, E. T. & Zhang, C. Z. Mechanisms of tandem duplication in the cancer genome. DNA Repair (Amst). 145, (2025).

7. Foo, T. K. & Xia, B. BRCA1-Dependent and Independent Recruitment of PALB2–BRCA2–RAD51 in the DNA Damage Response and Cancer. Cancer Res. 82, 3191–3197 (2022).

8. Sokolova, A., Johnstone, K. J., McCart Reed, A. E., Simpson, P. T. & Lakhani, S. R. Hereditary breast cancer: syndromes, tumour pathology and molecular testing. Histopathology 82, 70–82 (2023).

9. Lakhani, S. R. et al. Multifactorial analysis of differences between sporadic breast cancers and cancers involving BRCA1 and BRCA2 mutations. J. Natl. Cancer Inst. 90, 1138–1145 (1998).

10. Nik-Zainal, S. et al. Landscape of somatic mutations in 560 breast cancer whole-genome sequences. Nature 534, 47–54 (2016).

11. Menghi, F. et al. The tandem duplicator phenotype as a distinct genomic configuration in cancer. Proc. Natl. Acad. Sci. U. S. A. 113, E2373–E2382 (2016).

12. Yesbolatova, A. et al. The auxin-inducible degron 2 technology provides sharp degradation control in yeast, mammalian cells, and mice. Nat. Commun. 11, 5701 (2020).

13. Domingo, J., Baeza-Centurion, P. & Lehner, B. The Causes and Consequences of Genetic Interactions (Epistasis). Annu. Rev. Genomics Hum. Genet. 20, 433–460 (2019).

14. Lyu, X. et al. Human CST complex protects stalled replication forks by directly blocking MRE11 degradation of nascent-strand DNA. EMBO J. 40, (2021).

15. Barazas, M. et al. The CST Complex Mediates End Protection at Double-Strand Breaks and Promotes PARP Inhibitor Sensitivity in BRCA1-Deficient Cells. Cell Rep. 23, 2107–2118 (2018).

16. Mirman, Z. et al. 53BP1-RIF1-shieldin counteracts DSB resection through CST- and Polα-dependent fill-in. Nature 560, 112–116 (2018).

17. Chen, Y. H. et al. Interplay between the Smc5/6 complex and the Mph1 helicase in recombinational repair. Proc. Natl. Acad. Sci. U. S. A. 106, 21252–21257 (2009).

18. Xue, X. et al. Restriction of replication fork regression activities by a conserved SMC complex. Mol. Cell 56, 436–445 (2014).

19. Lambert, J. P. et al. Defining the budding yeast chromatin-associated interactome. Mol. Syst. Biol. 6, 448 (2010).

20. Bass, T. E. et al. ETAA1 acts at stalled replication forks to maintain genome integrity. Nat. Cell Biol. 18, 1185–1195 (2016).

21. Meetei, A. R. et al. A human ortholog of archaeal DNA repair protein Hef is defective in Fanconi anemia complementation group M. Nat. Genet. 37, 958–963 (2005).

22. Lai, X. et al. MUS81 nuclease activity is essential for replication stress tolerance and chromosome segregation in BRCA2-deficient cells. Nat. Commun. 2017 81 8, 1–13 (2017).

23. Feng, W. & Jasin, M. BRCA2 suppresses replication stress-induced mitotic and G1 abnormalities through homologous recombination. Nat. Commun. 2017 81 8, 1–15 (2017).

24. Blackford, A. N. et al. The DNA translocase activity of FANCM protects stalled replication forks. Hum. Mol. Genet. 21, 2005–2016 (2012).

25. Feng, S. et al. Profound synthetic lethality between SMARCAL1 and FANCM. Mol. Cell 84, 4522–4537.e7 (2024).

26. Fielden, J. et al. Comprehensive interrogation of synthetic lethality in the DNA damage response. Nature 640, (2025).

27. Zhang, S., Yu, Q., Li, Z., Zhao, Y. & Sun, Y. Protein neddylation and its role in health and diseases. Signal Transduct. Target. Ther. 2024 91 9, 1–36 (2024).

28. Basbous, J. & Constantinou, A. A tumor suppressive DNA translocase named FANCM. Crit. Rev. Biochem. Mol. Biol. 54, 27–40 (2019).

29. Blackford, A. N. et al. The DNA translocase activity of FANCM protects stalled replication forks. Hum. Mol. Genet. 21, 2005–2016 (2012).

30. Panday, A. et al. FANCM regulates repair pathway choice at stalled replication forks. Mol. Cell 81, 2428–2444.e6 (2021).

31. Deans, A. J. & West, S. C. FANCM Connects the Genome Instability Disorders Bloom’s Syndrome and Fanconi Anemia. Mol. Cell 36, 943–953 (2009).

32. Singh, T. R. et al. BLAP18/RMI2, a novel OB-fold-containing protein, is an essential component of the Bloom helicase–double Holliday junction dissolvasome. Genes Dev. 22, 2856 (2008).

33. Pierce, A. J., Johnson, R. D., Thompson, L. H. & Jasin, M. XRCC3 promotes homology-directed repair of DNA damage in mammalian cells. Genes Dev. 13, 2633–2638 (1999).

34. Mijic, S. et al. Replication fork reversal triggers fork degradation in BRCA2-defective cells. Nat. Commun. 2017 81 8, 1–11 (2017).

35. Schlacher, K. et al. Double-strand break repair-independent role for BRCA2 in blocking stalled replication fork degradation by MRE11. Cell 145, 529–542 (2011).

36. Pan, X. et al. FANCM, BRCA1, and BLM cooperatively resolve the replication stress at the ALT telomeres. Proc. Natl. Acad. Sci. U. S. A. 114, E5940–E5949 (2017).

37. Nacson, J. et al. BRCA1 Mutational Complementation Induces Synthetic Viability. Mol. Cell 78, 951–959.e6 (2020).

38. Pulver, E. M. et al. A BRCA1 coiled-coil domain variant disrupting PALB2 interaction promotes the development of mammary tumors and confers a targetable defect in homologous recombination repair. Cancer Res. 81, 6171 (2021).

39. Adam, S. et al. The CIP2A-TOPBP1 axis safeguards chromosome stability and is a synthetic lethal target for BRCA-mutated cancer. *Nat*. cancer 2, 1357–1371 (2021).

40. Haider, S. et al. The transcriptomic architecture of common cancers reflects synthetic lethal interactions. Nat. Genet. 2025 573 57, 522–529 (2025).

41. Ceccaldi, R. et al. Homologous-recombination-deficient tumours are dependent on Polθ-mediated repair. Nat. 2015 5187538 518, 258–262 (2015).

42. Mateos-Gomez, P. A. et al. Mammalian polymerase θ promotes alternative NHEJ and suppresses recombination. Nat. 2015 5187538 518, 254–257 (2015).

43. Krais, J. J. et al. Genetic separation of Brca1 functions reveal mutation-dependent Polθ vulnerabilities. Nat. Commun. 14, (2023).

44. Lim, K. S. et al. USP1 Is Required for Replication Fork Protection in BRCA1-Deficient Tumors. Mol. Cell 72, 925–941.e4 (2018).

45. Cadzow, L. et al. The USP1 Inhibitor KSQ-4279 Overcomes PARP Inhibitor Resistance in Homologous Recombination–Deficient Tumors. Cancer Res. 84, 3419 (2024).

46. Simoneau, A. et al. Characterization of TNG348: A Selective, Allosteric USP1 Inhibitor That Synergizes with PARP Inhibitors in Tumors with Homologous Recombination Deficiency. Mol. Cancer Ther. 24, 678–691 (2025).

47. Torrado, C., Ashton, N. W., D’Andrea, A. D. & Yap, T. A. USP1 inhibition: A journey from target discovery to clinical translation. Pharmacol. Ther. 271, (2025).

48. Cao, L. et al. A Selective Requirement for 53BP1 in the Biological Response to Genomic Instability Induced by Brca1 Deficiency. Mol. Cell 35, 534–541 (2009).

49. Bunting, S. F. et al. 53BP1 Inhibits Homologous Recombination in Brca1-Deficient Cells by Blocking Resection of DNA Breaks. Cell 141, 243–254 (2010).

50. Bouwman, P. et al. 53BP1 loss rescues BRCA1 deficiency and is associated with triple-negative and BRCA-mutated breast cancers. Nat. Struct. Mol. Biol. 17, 688–695 (2010).

51. Gupta, R. et al. DNA Repair Network Analysis Reveals Shieldin as a Key Regulator of NHEJ and PARP Inhibitor Sensitivity. Cell 173, 972–988.e23 (2018).

52. Noordermeer, S. M. et al. The shieldin complex mediates 53BP1-dependent DNA repair. Nature 560, 117–121 (2018).

53. Dev, H. et al. Shieldin complex promotes DNA end-joining and counters homologous recombination in BRCA1-null cells. Nat. Cell Biol. 2018 208 20, 954–965 (2018).

54. He, Y. J. et al. DYNLL1 binds to MRE11 to limit DNA end resection in BRCA1-deficient cells. Nat. 2018 5637732 563, 522–526 (2018).

55. Swift, M. L. et al. Dynamics of the DYNLL1–MRE11 complex regulate DNA end resection and recruitment of Shieldin to DSBs. Nat. Struct. Mol. Biol. 30, 1456–1467 (2023).

56. Tkáč, J. et al. HELB Is a Feedback Inhibitor of DNA End Resection. Mol. Cell 61, 405–418 (2016).

57. Hormeno, S. et al. Human HELB is a processive motor protein that catalyzes RPA clearance from single-stranded DNA. Proc. Natl. Acad. Sci. U. S. A. 119, (2022).

58. Chaudhuri, A. R. et al. Replication fork stability confers chemoresistance in BRCA-deficient cells. Nat. 2016 5357612 535, 382–387 (2016).

59. Taglialatela, A. et al. Restoration of Replication Fork Stability in BRCA1- and BRCA2-Deficient Cells by Inactivation of SNF2-Family Fork Remodelers. Mol. Cell 68, 414–430.e8 (2017).

60. Gari, K., Décaillet, C., Delannoy, M., Wu, L. & Constantinou, A. Remodeling of DNA replication structures by the branch point translocase FANCM. Proc. Natl. Acad. Sci. U. S. A. 105, 16107–16112 (2008).

61. Bizard, A. H. & Hickson, I. D. The dissolution of double Holliday junctions. Cold Spring Harb. Perspect. Biol. 6, (2014).

62. Harami, G. M. et al. The toposiomerase IIIalpha-RMI1-RMI2 complex orients human Bloom’s syndrome helicase for efficient disruption of D-loops. Nat. Commun. 13, (2022).

63. Scully, R., Elango, R., Panday, A. & Willis, N. A. Recombination and restart at blocked replication forks. Curr. Opin. Genet. Dev. 71, 154–162 (2021).

64. Servant, N., Allain, F., phupe, mdeloger & Philippe, L. R. bioinfo-pf-curie/nf-CRISPR: v1.0.0. doi:10.5281/ZENODO.8343750

65. Horlbeck, M. A. et al. Compact and highly active next-generation libraries for CRISPR-mediated gene repression and activation. Elife 5, (2016).

66. Llorca-Cardenosa, M. J. et al. SMG8/SMG9 Heterodimer Loss Modulates SMG1 Kinase to Drive ATR Inhibitor Resistance. Cancer Res. 82, 3962–3973 (2022).

## REFERENCES

1. Mengwasser, K. E. et al. Genetic Screens Reveal FEN1 and APEX2 as BRCA2 Synthetic Lethal Targets. Mol. Cell 73, 885–899.e6 (2019).

2. Srivastava, M. et al. HMCES safeguards replication from oxidative stress and ensures error-free repair. EMBO Rep. 21, (2020).

3. Álvarez-Quilón, A. et al. Endogenous DNA 3′ Blocks Are Vulnerabilities for BRCA1 and BRCA2 Deficiency and Are Reversed by the APE2 Nuclease. Mol. Cell 78, 1152–1165.e8 (2020).

4. Fleury, H. et al. The APE2 nuclease is essential for DNA double-strand break repair by microhomology-mediated end joining. Mol. Cell 83, 1429–1445.e8 (2023).

5. Farmer, H. et al. Targeting the DNA repair defect in BRCA mutant cells as a therapeutic strategy. Nature 434, 917–921 (2005).

6. Bryant, H. E. et al. Specific killing of BRCA2-deficient tumours with inhibitors of poly(ADP-ribose) polymerase. Nature 434, 913–917 (2005).

7. Zimmermann, M. et al. CRISPR screens identify genomic ribonucleotides as a source of PARP-trapping lesions. Nature 559, 285 (2018).

8. Adam, S. et al. The CIP2A-TOPBP1 axis safeguards chromosome stability and is a synthetic lethal target for BRCA-mutated cancer. *Nat*. cancer 2, 1357–1371 (2021).

9. Jensen, R. B., Ozes, A., Kim, T., Estep, A. & Kowalczykowski, S. C. BRCA2 is epistatic to the RAD51 paralogs in response to DNA damage. DNA Repair (Amst). 12, 306 (2013).

10. Qing, Y. et al. The Epistatic Relationship between BRCA2 and the Other RAD51 Mediators in Homologous Recombination. PLOS Genet. 7, e1002148 (2011).

11. van de Kooij, B. et al. EXO1 protects BRCA1-deficient cells against toxic DNA lesions. Mol. Cell 84, 659–674.e7 (2024).

12. Ludwig, T., Chapman, D. L., Papaioannou, V. E. & Efstratiadis, A. Targeted mutations of breast cancer susceptibility gene homologs in mice: lethal phenotypes of Brca1, Brca2, Brca1/Brca2, Brca1/p53, and Brca2/p53 nullizygous embryos. Genes Dev. 11, 1226–1241 (1997).

13. Gasco, M., Yulug, I. G. & Crook, T. TP53 mutations in familial breast cancer: Functional aspects. Hum. Mutat. 21, 301–306 (2003).

14. Tkáč, J. et al. HELB Is a Feedback Inhibitor of DNA End Resection. Mol. Cell 61, 405–418 (2016).

15. He, Y. J. et al. DYNLL1 binds to MRE11 to limit DNA end resection in BRCA1-deficient cells. Nat. 2018 5637732 563, 522–526 (2018).

